# Partial metal ion saturation of C2 domains primes Syt1-membrane interactions

**DOI:** 10.1101/810010

**Authors:** Sachin Katti, Sarah B. Nyenhuis, Bin Her, David S. Cafiso, Tatyana I. Igumenova

## Abstract

Synaptotagmin 1 (Syt1) is an integral membrane protein that acts as a Ca^2+^ sensor of neurotransmitter release. How the Ca^2+^-sensing function of Syt1 is coupled to its interactions with anionic membranes and synaptic fusion machinery is not well understood. Here, we investigated the dynamics and membrane-binding properties of Syt1 under conditions where its highest affinity Ca^2+^ sites, which are thought to drive the initial membrane recruitment, are selectively populated by divalent metal ions. To create such protein states for the Ca^2+^-sensing C2 domains of Syt1, we exploited the unique chemistry of Pb^2+^, a xenobiotic metal ion that is isostructural and isofunctional to Ca^2+^. NMR experiments revealed that binding of a single metal ion results in the loss of conformational plasticity of the C2 domain loop regions that are involved in both coordinating metal ions and membrane interactions. In the C2A domain, a single metal ion is sufficient to drive its weak association with PtdSer-containing membranes; in C2B, it enhances the interactions with the signaling lipid PtdIns(4,5)P_2_. In full-length Syt1, both C2 domains associate with PtdSer-containing membranes, with the depth of insertion modulated by the occupancy of the metal ion sites. Our data suggest that Syt1 adopts a shallow membrane-bound state upon initial recruitment of its C2 domains to the membranes. The properties of this state, such as conformationally restricted loop regions and positioning of C2 domains in close proximity to anionic lipid headgroups, “prime” Syt1 for binding a full complement of metal ions required for activation of protein function.

---

The process of neurotransmitter release is tightly coupled to the changes in neuronal Ca^2+^ levels (1). Synaptotagmin 1 (Syt1), an integral membrane protein that is anchored to synaptic vesicles through its N-terminal region, plays a major regulatory role in this process (2–4). Together with its protein effectors, such as SNAREs (soluble N-ethylmaleimide sensitive factor attachment protein receptors) (5,6) and complexin (7), Syt1 initiates the vesicle fusion in a Ca^2+^-dependent manner (8). This results in the opening of membrane fusion pore, through which neurotransmitters are released from the vesicles into the synaptic cleft (9,10).

The Ca^2+^-sensing function of Syt1 resides on its cytosolic C-terminal region that comprises tandem C2A and C2B domains, connected by a 9-residue linker. Several models have been proposed to explain the membrane-binding modes of these C2 domains and their contributions to the overall Ca^2+^ response (11–15). However, the exact relationship between variations in cytosolic Ca^2+^ levels and the C2-membrane interactions remains elusive, since most studies were conducted at supra-physiological Ca^2+^ concentrations.

The objective of this work was to gain a mechanistic insight into the membrane recruitment of Syt1 C2 domains, under conditions that mimic low Ca^2+^ concentrations. This regime is relevant to two aspects of Syt1 function: regulation of spontaneous neurotransmitter release that occurs at stochastic Ca^2+^ elevations in the absence of action potentials (16); and the initial stages of C2-membrane recruitment that would involve interactions with cytosolic Ca^2+^ prior to the binding to negatively charged lipids.

The C2 domains of Syt1 together have five Ca^2+^-binding sites: three in C2A and two in C2B. These sites reside on the aspartate-rich apical loop regions of the C2 domains (17,18). Ca^2+^ binding facilitates the association of C2 domains with anionic phospholipids, such as phosphatidylserine (PtdSer) and phosphatidylinositol-(4,5)-bisphosphate (11,19). The role of metal ions in the membrane association process is thought to be two-fold: (i) to make the C2 loops more electropositive and thereby enhance the Coulombic attraction between the protein and anionic lipids; and (ii) to directly coordinate anionic phospholipids.

In each C2 domain, the differences between Ca^2+^ affinities of the first two sites are within one order of magnitude (17,18). Site 1, shown in **Fig. 1A** in the context of Pb^2+^-complexed C2 domains (*vide infra*), has the highest Ca^2+^ affinity and would be expected to be primarily populated at low-to moderate concentrations of Ca^2+^. In fact, in another C2-containing protein, protein kinase Cα, population of Site 1 by Ca^2+^ was proposed to be responsible for the initial recruitment of the protein to membranes (20). However, the single Ca^2+^-bound states of C2 domains are inaccessible for structural and dynamic characterization, because comparable Ca^2+^ affinities result in the formation of multiple metal ion-bound C2 domain species in solution.

**Figure 1.**
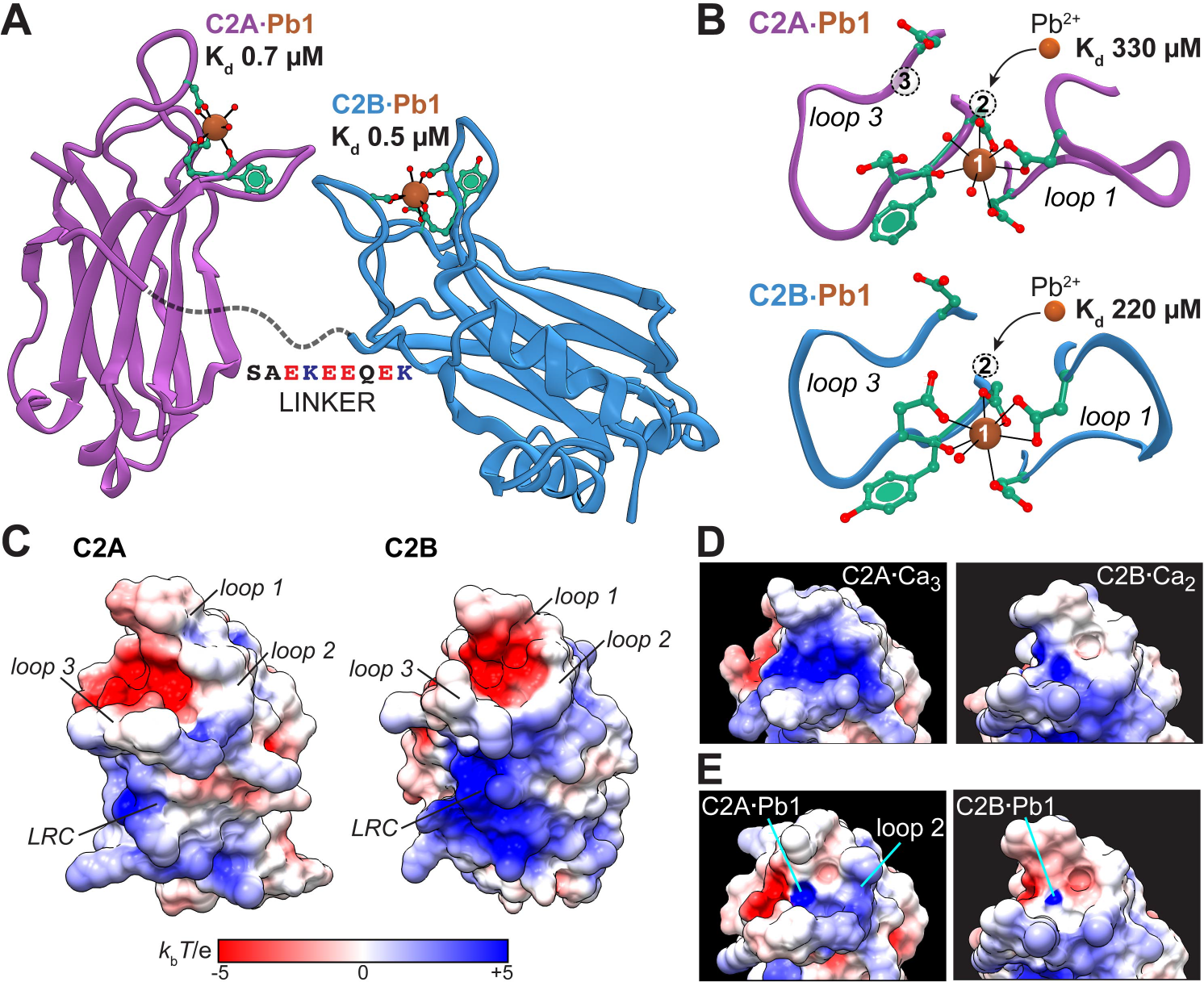
High-affinity Pb^2+^-binding sites of Syt1. (**A**) Cartoon representation of the Syt1 cytoplasmic region assembled from the Pb^2+^-bound crystal structures of isolated C2 domains, C2A (5VFE) and C2B (5VFG). The Pb^2+^ affinities to Site 1 exceed those of Ca^2+^ by ~2 orders of magnitude. (**B**) Expansion of loop regions in C2A·Pb1 and C2B·Pb1 showing the numbered metal-ion binding sites and the low affinity of Pb^2+^ to Site 2. (**C**) Electrostatic potential maps of apo C2A and C2B. (**D,E**) Electrostatic potential maps of loop regions with full complement of metal ions (**D**) and with Site 1 selectively populated by Pb^2+^ (**E**). The maps were generated using Adaptive Poisson- Boltzmann Solver (APBS) plugin of UCSF Chimera. The PDB IDs of structures used: 4WEE (C2A apo), 1BYN (C2A·Ca_3_), 5CCJ (C2B apo), and 1TJX (C2B·Ca_2_).

To overcome this challenge and create a dominant population of single metal-bound C2 species, we made use of Pb^2+^, a Ca^2+^-mimicking xenobiotic cation (21). Heavy metal ions were shown to be useful tools to establish the C2 structure-function relationship (22–25). What sets Pb^2+^ apart from other metal ions is two features. First, it supports the membrane interactions of C2 domains and is therefore isofunctional to Ca^2+^ (24,26,27). Second, although Pb^2+^ populates the exact same C2 domain sites as Ca^2+^ in solution, there are significant differences in its affinities to Site 1 and 2-3: the affinity of Pb^2+^ to Site 1 is ~450-fold higher than that to Site 2 (**Fig. 1B**) (27). These unique properties of Pb^2+^ with respect to its interactions with C2 domains enabled us to simultaneously achieve selective population of specific metal binding site, Site 1, with minimum perturbation of the membrane binding function of the proteins.

For both C2 domains of Syt1, we found that population of Site 1 by a divalent metal ion causes rigidification of their membrane-binding regions. Using isotropically tumbling bicelles and large unilamellar vesicles as membrane mimics, we characterized the membrane-binding properties of Pb^2+^-complexed C2 domains: individual, in tandem, and in the context of full-length Syt1. Our work provides evidence that Syt1 can be “primed” for productive membrane interactions by partial metal-ion saturation of loop regions.

## RESULTS

### Electrostatic properties of C2 domains in different states of metal ligation

We previously obtained high-resolution crystal structures of individual Pb^2+^-complexed Syt1 C2 domains, where Pb^2+^ ions bind to Site 1 with two-orders of magnitude higher affinity than Ca^2+^ (27) (**Figs. 1A,B**). To determine how metal ion binding to Site 1 alters the electrostatic profiles of these C2 domains, we calculated the surface electrostatic potential maps of the proteins in different states of metal ligation. The negatively charged metal ion-binding region is formed by the aspartate-rich loops (**Fig. 1C**). In the presence of a full complement of Ca^2+^ ions, C2A undergoes a prominent electrostatic shift where the intra-loop potential changes from negative to positive (**Fig. 1D**). The effect of bound Ca^2+^ on the same region in C2B is the neutralization of the negative charge (**Fig. 1D**). This is because C2A can accommodate 3 Ca^2+^ ions within the intra-loop region, as compared to the C2B’s maximum of 2. Pb^2+^ binding to Site 1 partially neutralizes the intra-loop region in both domains. Due to the differences in the arrangement of basic residues, loop 2 of C2A becomes more electropositive while loop 2 of C2B becomes neutral (**Fig. 1E**). The differences in electrostatic properties between the two C2 domains determine how the single metalion bound states interact with anionic membranes (*vide infra*).

### Population of Site 1 by a metal ion alters conformational plasticity of the membrane-binding regions

In addition to electrostatics, the conformation and dynamics of the membrane-binding regions in the C2·Pb1 complexes can have profound influence on the membrane association. The average conformation of the domains, as it exists in the crystalline state, changes only moderately upon metal ion binding. The ms-to-μs, but not the sub-nanosecond backbone dynamics in a related C2 domain were previously shown to be responsive to the state of metal ligation (28). We therefore investigated the ms-to-μs dynamics of the Syt1 C2 domains in apo and single-Pb^2+^ states using solution NMR methods.

Two types of NMR parameters, R_2,HE_ and R_2,CPMG_, were measured for all spectrally resolved N-H groups of the C2 domain backbone. R_2,HE_ is a free-precession transverse relaxation rate constant, whose elevated values are indicative of ms-to-μs conformational exchange processes. R_2,CPMG_ is a transverse relaxation rate constant whose elevated values reflect dynamics on timescales faster than 100 μs. A comparison of the two transverse relaxation rate constants enables one to estimate, in a residue-specific manner, the timescale of motions present in macromolecules.

The NMR relaxation data for the apo C2 domains are presented in **Fig. 2**. Based on the elevated R_2,HE_ values (**Fig. 2A**), the backbone of apo C2A has two highly dynamic regions: loop 3, which participates in both metal-ion coordination and binding to PtdSer-rich membranes; and the lysine-rich cluster (LRC). Surprisingly, there is little dynamics in loop 1, which provides roughly half of the oxygen ligands when Site 1 gets populated by a divalent metal ion.

**Figure 2.**
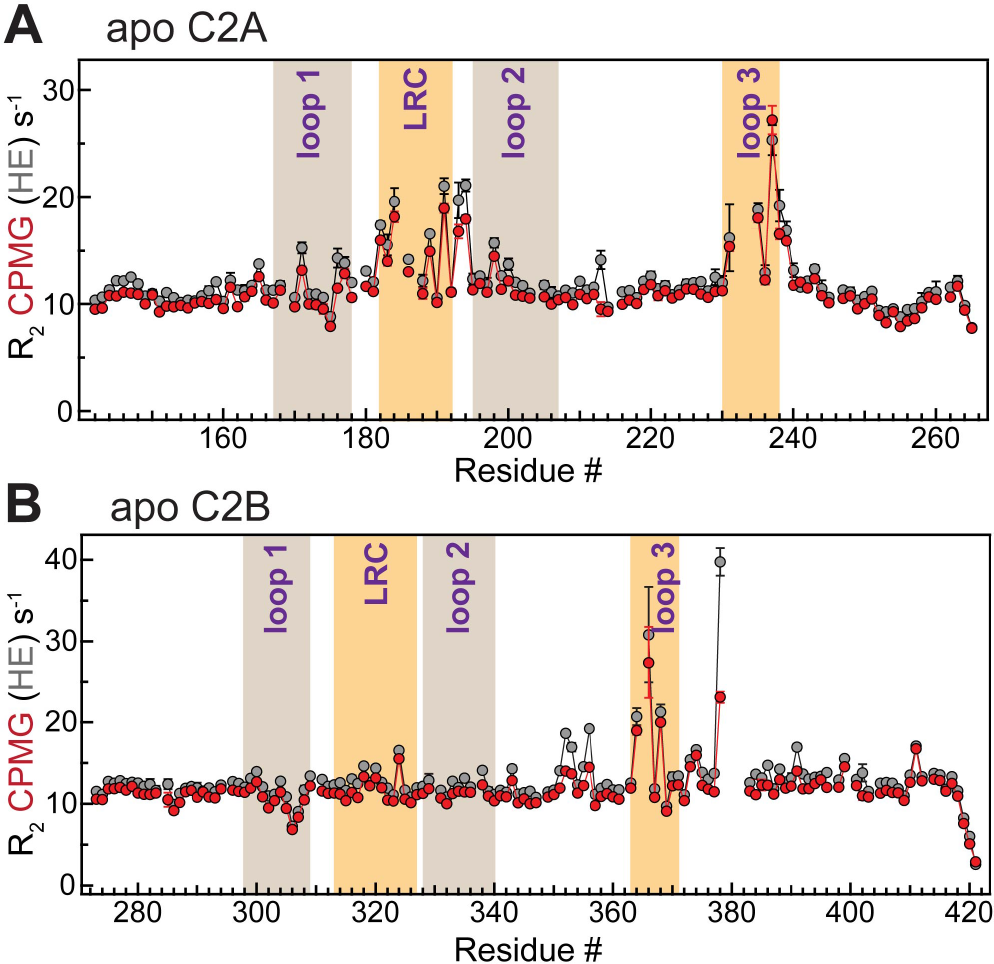
The membrane-binding regions of apo C2A and C2B are dynamic on the microsecond timescale. R_2,HE_ (gray) and R_2,CPMG_ (red) are plotted against the amino acid sequence of C2A (**A**) and C2B (**B**). The regions corresponding to metal-ion coordinating loops and lysine-rich clusters are highlighted.

The dynamics profile is different for C2B (**Fig. 2B**). The most significant difference between C2B and C2A is located in the LRC that shows no conformational exchange in C2B but is highly dynamic in C2A. The functional role of LRC in C2B domains is to provide an additional docking site for the negatively charged signaling lipids, such as PtdIns(4,5)P_2_, and thereby increase the residence time at the membrane (29). Another noteworthy difference is that although in both domains loop 3 undergoes conformational exchange, only in C2B the regions adjacent to loop 3 hinges are also dynamic. To evaluate the timescale, we inspected the difference plots between the R_2,HE_ and R_2,CPMG_ values. In both domains, the attenuation of dynamics due to the application of the CPMG pulse train is modest (**Fig. S1A** and **S1B**). These data indicate that the timescale of loop 3 and LRC (in C2A) motions is faster than 100 μs.

To determine how metal ion binding to Site 1 influences the backbone dynamics, we prepared single-Pb^2+^ complexes of the C2 domains and measured their R_2,HE_ and R_2,CPMG_ values. Under 1:1 stoichiometric conditions of our sample, Site 1 is slightly under-populated. Although the population of apo proteins is very low (~2-3 %), the inter-conversion between apo and metal-ion bound species can lead to large chemical exchange effects on the millisecond timescale, as was previously shown for the Ca^2+^ ion binding to the C2A variant (30). Indeed, we observed a significant elevation of the R_2,HE_ values in the loop regions of both domains. Fortunately, these processes occur on a timescale of ms and therefore can be efficiently suppressed by an application of the CPMG pulse train (**Fig. S2**). Therefore, the R_2,CPMG_ values in the C2·Pb1 complexes report almost exclusively on the intrinsic dynamics of the loops.

The R_2,CPMG_ differences between the Pb^2+^-complexed and apo proteins revealed that metal ion binding to Site 1 attenuates the dynamics of loop 3 (**Fig. 3**). This is manifested in large negative values of ΔR_2,CPMG_ in loop 3 for both domains. In addition, the entire positively charged LRC region of C2A looses its conformational plasticity (**Fig. 3A**). This indicates that the dynamic effect of metal ion binding to Site 1 propagates as far as ~15-25 Å throughout the C2A domain in what could be an allosteric communication between the metal ion binding sites and LRC. Loop 1, which is immobile on the ms-to-μs timescales in the apo-states of C2A and C2B (**Fig. 2**), experiences a slight increase in R_2,CPMG_ due to incomplete suppression of the ms-timescale chemical exchange due to metal ion binding (*vide supra*).

**Figure 3.**
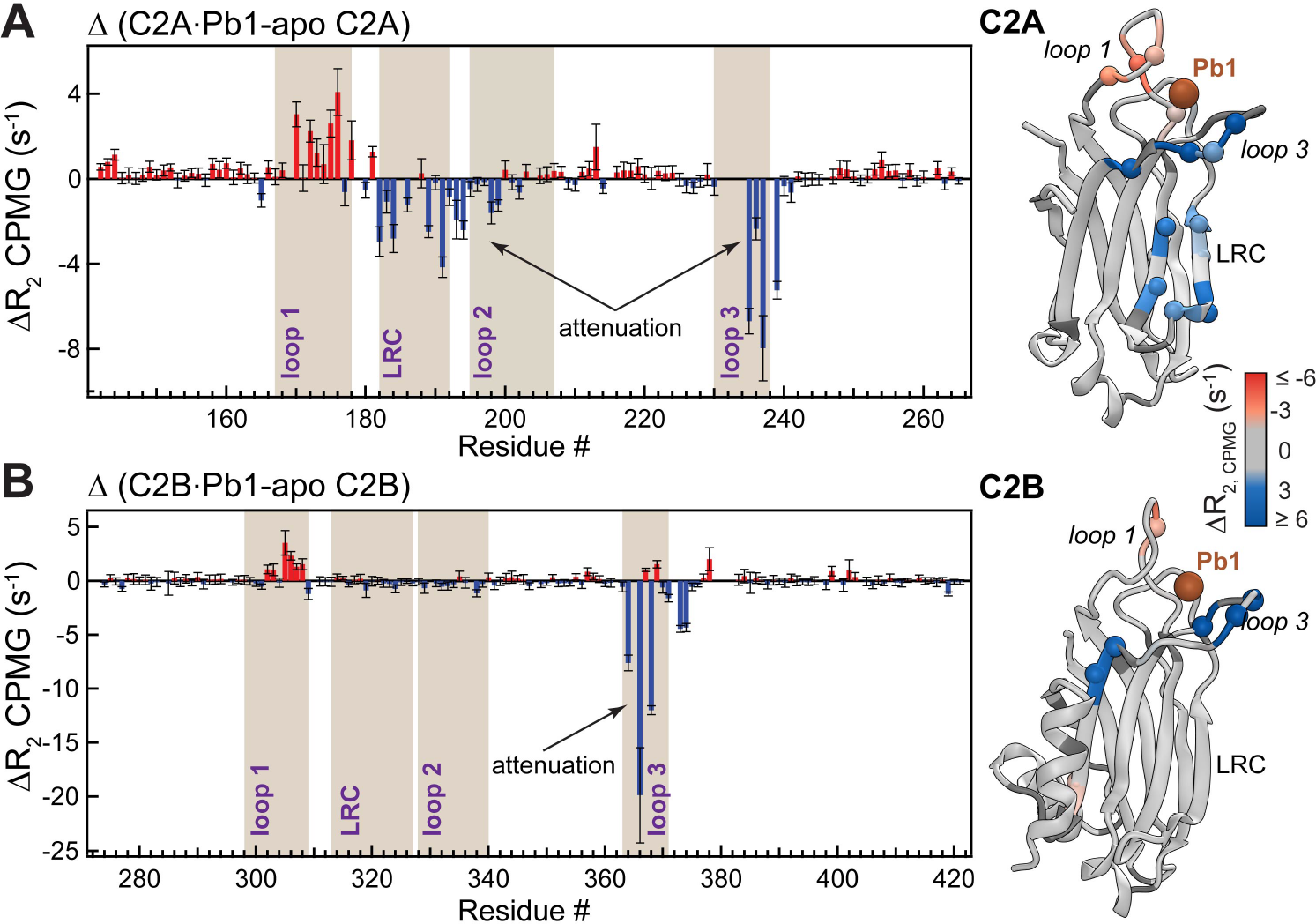
Binding of a single metal ion to C2 attenuates the μs-timescale dynamics of the membrane-binding regions. Difference plots of the R_2,CPMG_ values measured in apo and Pb^2+^-complexed C2 domains are shown for the C2A (**A**) and C2B (**B**) domains. Negative values indicate the attenuation of dynamics. The R_2,CPMG_ differences are color-coded and mapped onto the 3D structures of the Pb^2+^-complexed C2 domains.

For the function of C2 domains, the implications of the attenuated dynamics are two-fold. First, conformational restriction brought about by a metal ion in Site 1 may facilitate membrane interactions through imposing loop 3 orientation that is favorable for membrane insertion. Second, the geometric restriction of the Asp ligands, some of which are shared by metal ions in Sites 1 and 2, will likely facilitate the subsequent metal-ion binding to Site 2. In either case, the effect of populating Site 1 with a metal ion would “prime” the C2 domains of Syt1 for metal-ion dependent membrane interactions.

### C2A weakly associates with anionic membranes in a single metal-ion bound state

We then asked if the population of Site 1 by a metal ion is sufficient to support the membrane interactions of the Syt1 C2 domains. Starting with C2A, we prepared the [U-^15^N]-enriched C2A·Pb1 complex and mixed it with the preparation of isotropically tumbling bicelles that mimic the membrane environment. Two anionic phospholipids: phosphatidylserine (PtdSer) and phosphatidylinositol-4,5-bisphosphate (PtdIns(4,5)P_2_) were incorporated into the bicelles at 14% and 2%, respectively, relative to the long-chain phospholipid. Comparison of the [^15^N-^1^H] HSQC spectra with and without bicelles (**Fig. 4A**) revealed two spectroscopic signatures: chemical shift perturbations (CSPs, Δ) of the N-H cross-peaks and attenuation of their intensities. The CSPs affected loop regions 1, 2, and 3 of the C2A domain, but not the lysine-rich cluster (**Figs. 4B,D**). No chemical shift perturbations were observed when C2A domain in the absence of any metal ions was mixed with bicelles, indicating that CSPs report specifically on the metal-ion dependent interactions of the C2A domain with membranes. Lack of CSPs in the LRC indicates that this region does not directly interact with PtdSer and PtdIns(4,5)P_2_. In the membrane-free system, a similar interaction pattern was observed between the Ca^2+^-complexed C2A domain and inositol-1,4,5-triphosphate (29).

**Figure 4.**
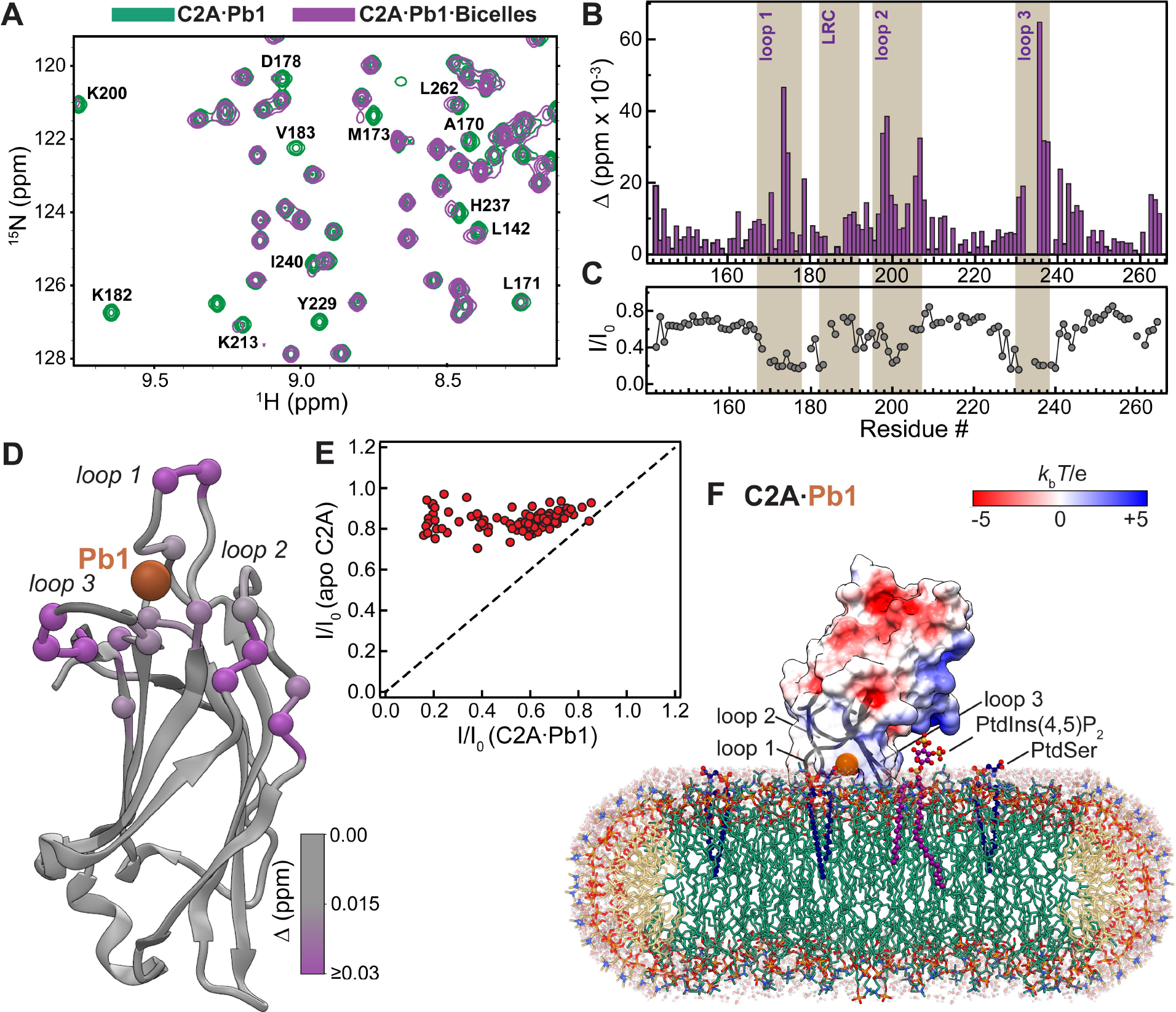
Syt1 C2A·Pb1 complex interacts with anionic bicelles. (**A**) Overlay of the [^15^N-^1^H] HSQC spectral regions of C2A·Pb1 in the absence (green) and presence (purple) of anionic bicelles that illustrates CSPs experienced and broadening of protein resonances upon interactions with bicelles. (**B**) CSP plot showing that perturbations are localized primarily to the loop regions. (**C**) Attenuation of peak intensities in the C2A·Pb1 complex upon addition of anionic bicelles, expressed as residue-specific I/I_0_ ratios, where I and I_0_ are the peak intensities in the presence and absence of bicelles, respectively. Loop regions are preferentially broadened due to chemical exchange. (**D**) CSP values mapped on the C2A·Pb1 complex (PDB ID: 5VFE) illustrating the proximity of the affected regions to the metal ion binding Site 1. (**E**) Comparison of the residue-specific I/I_0_ ratios in apo C2A and C2A·Pb1. The overall intensity attenuation is significant in the C2A·Pb1 sample, but negligible in the apo C2A sample, where it merely reflects an increase in solution viscosity due to bicelle addition. (**F**) Schematic representation of the C2A·Pb1-bicelle complex that is consistent with our experimental data.

The intensity attenuation of the C2A·Pb1 N-H cross-peaks, expressed as the ratio of intensities in the bicelle-containing (I) and bicelle-free (I_0_) samples varied across the primary structure (**Fig. 4C**). The regions with the most reduced intensities correspond to the protein loops and mirror the CSP pattern. This is a manifestation of a chemical exchange process whose kinetics is in the intermediate-to-fast regime on the NMR chemical shift timescale. It is well established that the PtdSer can coordinate the C2-bound metal ions while interacting simultaneously with the loop residues (31,32). We conclude that the chemical exchange process represents the binding of the binary C2A·Pb1 complex to the bicelles, with the formation of the ternary C2A·Pb1·PtdSer(bicelle) complex. This conclusion is further reinforced by the comparison of the residue-specific I/I_0_ ratios between apo C2A and C2A·Pb1 in the presence of bicelles (**Fig. 4E**). The intensity ratios are systematically lower for the C2A·Pb1 complex, and not just for the loop regions. The overall decrease in intensities is due to the increase in the apparent rotational correlation time of C2A·Pb1, because it partially associates with bicelles that are slow-tumbling high-molecular-mass entities.

The C2A·Pb1 association with membranes also occurs in the absence of PtdIns(4,5)P_2_, where the only anionic lipid component is PtdSer. We demonstrated this by conducting NMR-detected experiments of the C2A·Pb1 binding to large unilamellar vesicles (**Fig. S3**). Moreover, the fraction of metal-complexed C2A bound to membranes increases approximately 2-fold when, in addition to Site 1, Site 2 gets populated by Pb^2+^ (**Fig. S4**).

Taken together, our data indicate that population of Site 1 by a divalent metal ion is sufficient to drive weak association of the C2A domain with anionic membranes. The membrane-interacting regions primarily involve the metal-binding loops, as shown schematically for the C2·Pb1·bicelle complex in **Fig. 4E**.

### C2B interactions with anionic membranes are enhanced by metal ion binding to Site 1

The C2B domain in Syt1 is distinct from C2A with respect to its electrostatic properties. The unique electrostatic makeup of the C2B surface provides additional complexity and control to the membrane interactions of Syt1 (15,18,33,34). The LRC of C2B has 6 basic residues while that of C2A has 4 basic residues that are flanked by two acidic ones. This makes the LRC of C2B significantly more electropositive (see **Fig. 1C**), turning it into an effective PtdIns(4,5)P_2_ sensor (35). Moreover, there are additional basic residues, Arg 398 and Arg 399, at the C2B end opposite to the loop region. These residues are implicated in the membrane-apposition process (12,34). In the presence of Ca^2+^, the basic regions can create scaffolds by simultaneously interacting with various lipid and protein partners such as PtdSer, inositol polyphosphates, Ca^2+^ channels, and SNAREs (15,36–38). In the absence of Ca^2+^, the LRC of C2B is believed to pre-associate with the PtdIns(4,5)P_2_-rich plasma membranes (39).

Using the same experimental strategy as for C2A, we prepared the [U-^15^N]-enriched C2B·Pb1 and collected 2D [^15^N-^1^H] HSQC NMR spectra in the presence and absence of anionic bicelles (**Fig. 5A**). Extensive line broadening accompanied by significant chemical shift changes was observed for many N-H cross-peaks of C2B. The CSP pattern calculated for the bicelle-containing samples relative to bicelle-free C2B·Pb1 was clearly distinct from what we observed for the C2A·Pb1 complex in two aspects: (i) the CSPs for loop 3 of C2B were significantly larger than those for loop 1, suggesting a more prominent role of loop 3 in protein-membrane interactions; and (ii) there were very significant CSPs in the LRC-loop 2 region (compare **Figs. 4B** and **5B**). Mapping the CSP values onto the 3D structure of the Pb^2+^-complexed C2B domain highlighted a contiguous surface likely to be involved in interactions with anionic membranes (**Fig. 5D**). Another drastic difference between C2A·Pb1 and C2B·Pb1 was in the extent of signal attenuation upon addition of bicelles (compare **Figs. 4C** and **5C**). The intensity drop for the C2B·Pb1 was uniform, with the exception of a few flexible C-terminal residues, and substantial: the mean I/I_0_ value from the 5% trimmed data set was 0.14. We conclude that the C2B·Pb1 complex, in contrast to C2A·Pb1, associates with anionic bicelles with high affinity.

**Figure 5.**
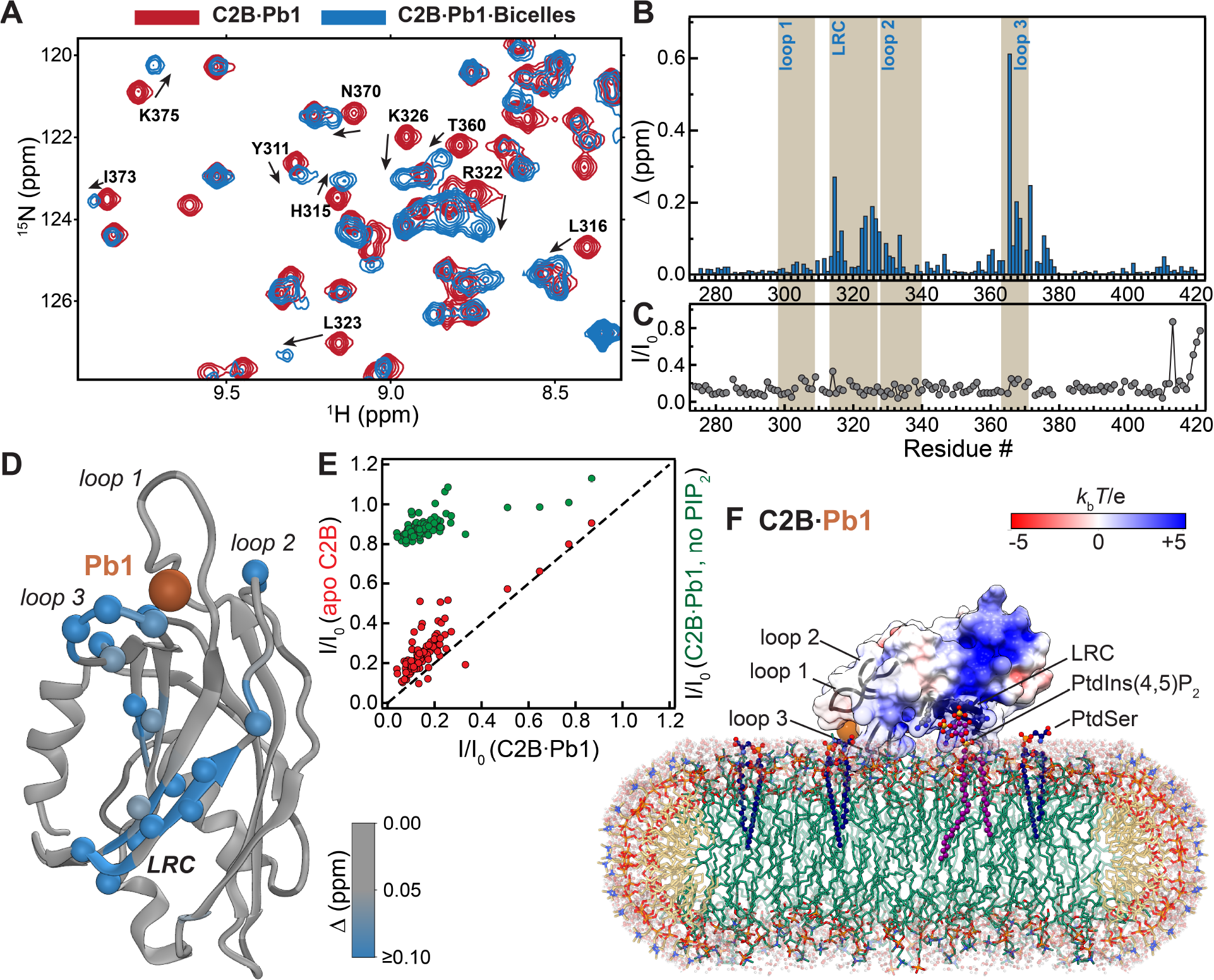
C2B·Pb1 binds to anionic bicelles with high affinity. (**A**) Overlay of the [^15^N-^1^H] HSQC spectral regions of C2B·Pb1 in the absence (red) and presence (blue) of anionic bicelles that illustrates CSPs and broadening of protein resonances upon interactions with bicelles. (**B**) CSP plot showing that perturbations are localized primarily to the LRC and loop 3 of the C2B domain. (**C**) Attenuation of peak intensities in the C2B·Pb1 complex upon addition of anionic bicelles, expressed as residue-specific I/I_0_ ratios, where I and I_0_ are the peak intensities in the presence and absence of bicelles, respectively. Significant uniform (with the exception of the flexible C-term residues) attenuation indicates high-affinity association of C2A·Pb1 with bicelles. (**D**) CSP values mapped on the C2B·Pb1 complex (PDB ID: 5VFG) cover a contiguous surface that includes primarily loop 3 and the LRC. (**E**) Correlation of residue-specific I/I_0_ ratios in apo C2B and C2B·Pb1 in the presence of PtdIns(4,5)P_2_ (red) indicates that metal ion binding to Site 1 contributes moderately to protein-bicelle interactions. Omitting PtdIns(4,5)P_2_ abolishes the interactions of C2B·Pb1 with bicelles (green). (**F**) Schematic representation of the C2B·Pb1-bicelle complex that is consistent with our experimental data.

To establish the metal-ion dependence of this process, we conducted a control experiment in which the metal-ion free C2B domain (apo C2B) was mixed with anionic bicelles of the same composition. The observed CSP values were smaller than those for the C2B·Pb1 complex, especially for loop 3 (**Fig. S5**). The correlation of I/I_0_ intensity ratios between apo C2B and C2B·Pb1 showed systematically lower values for the latter, suggesting that the fractional population of the membrane-bound C2B·Pb1 species is higher than that of the apo species (**Fig. 5E**, red symbols). However, the overall effect is rather modest, suggesting that the presence of metal ion at Site 1 contributes moderately to the C2B-membrane interactions. The binding of C2B·Pb1 to bicelles is significantly enhanced by PtdIns(4,5)P_2_: when we omitted PtdIns(4,5)P_2_, the I/I_0_ ratios increased to values of 0.8 and above (**Fig. 5E**, green symbols).

Our data are consistent with the orientation of C2B·Pb1 where the LRC and loop 3, but not loop 1, are in close membrane contact (**Fig. 5F**). In the previously reported model of the C2B-membrane complex under the saturating Ca^2+^ conditions (14,15), C2B has both loops 1 and 3 inserted into the membrane, while its LRC makes contact with PtdIns(4,5)P_2_. The implication is that the metal-ion binding to Site 2 likely causes insertion of loop 1 into the headgroup region and “tilts” the C2B by reducing the angle between its long axis and the bilayer normal.

### The affinity of C2B to PtdIns(4,5)P_2_ is enhanced by metal ion at Site 1

Binding of a full complement of Ca^2+^ ions enhances the affinity of C2 domains to PtdIns(4,5)P_2_ (19,26). To evaluate the thermodynamic gain afforded exclusively by Site 1, we conducted NMR-detected binding experiments between C2B (apo and the C2B·Pb1 complex) and di-C4-PtdIns(4,5)P_2_. The rationale behind using water-soluble di-C4-PtdIns(4,5)P_2_, as compared to inositol 1,4,5-trisphosphate, is that it has the same electrostatic properties of the headgroup as the longer-chain PtdIns(4,5)P_2_.

Both C2B species responded to increasing concentration of PtdIns(4,5)P_2_ in a titratable manner (**Fig. 6A** and **Fig. S6A**). The binding process is in the fast exchange regime relative to the NMR chemical shift timescale. This is manifested in a smooth trajectory of the N-H cross-peaks as a function of increasing concentration of ligand in both apo C2B and C2B·Pb1 [^15^N,^1^H] HSQC spectra. The fast exchange behavior enabled us to construct the binding curves that represent the chemical shift changes experienced by the N-H cross-peaks, Δ, as a function of total concentration of di-C4-PtdIns(4,5)P_2_. Binding curves for well-resolved residues that showed a combined Δ of > 0.05 ppm were globally fitted to obtain the following dissociation constants: K_d_ = 102 ± 3 μM (C2B·Pb1) and 215 ± 16 μM (apo C2B). We conclude that the presence of metal ion in Site 1 of the C2B domain alone is sufficient to cause 2-fold increase of affinity to PtdIns(4,5)P_2_. A comparison of the CSP patterns (**Fig. S6**) mapped onto the protein 3D structures (**Fig. 6**) shows that the same regions in apo C2B and the C2B·Pb1 are influenced by the PtdIns(4,5)P_2_ interactions: the LRC and loop 3-adjacent β7 segment (**Fig. 6B**). These regions form a concave site that is typical for C2 domains with specificity towards phosphoinositides (35,40).

**Figure 6.**
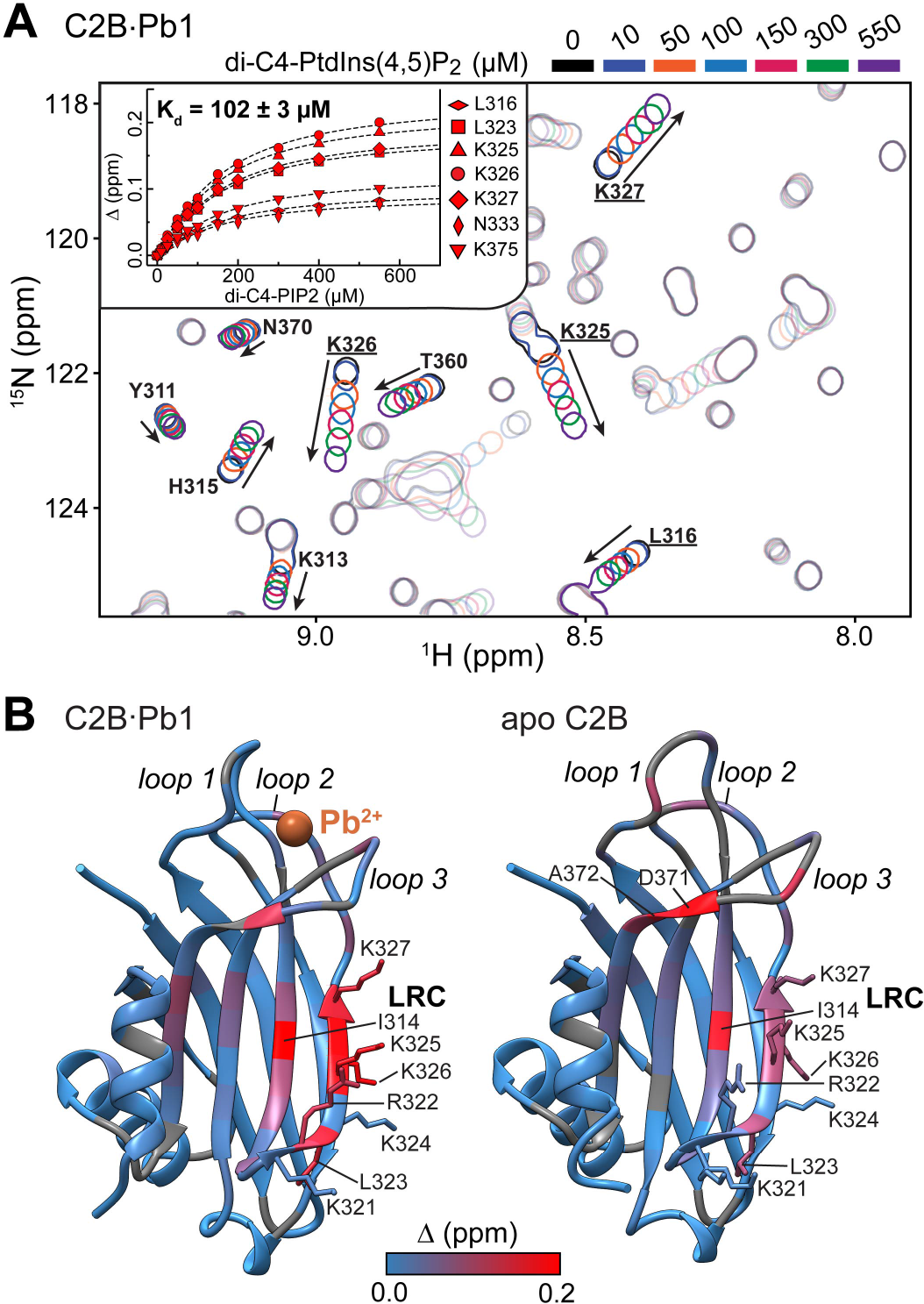
Binding of Pb^2+^ at Site 1 enhances the affinity of C2B towards PtdIns(4,5)P_2_. (**A**) Overlay of the [^15^N-^1^H] HSQC spectral regions showing progressive chemical shift changes upon addition of di-C4-PtdIns(4,5)P_2_ to the C2B·Pb1 complex. N-H cross-peaks that illustrate the binding of di-C4-PtdIns(4,5)P_2_ to the canonical site in the LRC are labeled. Inset: NMR-detected binding curves for residues that either form or are in close proximity to the canonical binding site. The effective K_d_ value extracted from the global fit of the curves is 102 μM, which is approximately 2-fold lower than that for the apo C2B. (**B**) Chemical shift perturbations Δ due to di-C4-PtdIns(4,5)P_2_ binding (0-0.4 mM range) mapped onto the crystal structures the C2B·Pb1 complex (5VFG) and apo C2B (5CCJ). Positively charged residues that form LRC are shown in stick representation. Additionally, residues experiencing Δ larger than 0.15 ppm are labeled.

The 2-fold increase in affinity represents the thermodynamic enhancement that is attributed solely to the interactions of the LRC residues with PtdIns(4,5)P_2_. In neuronal membranes, an additional favorable contribution to the protein-membrane interactions would come from higher-abundance anionic phospholipids, such as PtdSer, which directly coordinates the metal ion bound to Site 1 of C2B (31).

### The membrane interaction pattern of C2AB is similar to that of individual domains

The tandem C2A and C2B domains (**Fig. 1A**) constitute the membrane-binding unit of Syt1 (18,41). Their distinct electrostatic properties and preferences towards binding partners govern the Syt1-mediated vesicle fusion (15,42,43). The question of the C2AB conformational preferences has elicited some discussion in the literature. In crystalline state, C2AB can adopt a “closed” conformation with well-defined inter-domain interface (44). In solution, the atomic force microscopy and intramolecular FRET measurements by Chapman and coworkers indicate that there exists an interaction between C2A and C2B (45) and that it is important for the Syt1 function (46). However, NMR experiments conducted by Rizo’s group argue against the presence of a well-defined inter-domain interface in solution (12). What is evident is that there exists significant inter-domain conformational flexibility in the C2AB fragment. We sought to determine how this flexibility influences membrane interaction properties under limiting metal ion conditions.

Using the C2AB fragment, we first tested if Pb^2+^ populates the same metal-ion binding sites as it does in the isolated domains. Due to the acidic nature of the 9-residue linker (there are 4 glutamate residues, **Fig. 1A**), there was a possibility that this region could sequester added Pb^2+^. [^15^N-^1^H] TROSY HSQC spectra of [U-^15^N, ~80%-^2^H] C2AB at 1:1 and 1:2 protein-to-Pb^2+^ ratios (**Fig. S7**) revealed the same chemical exchange regime, saturation behavior, and chemical shift perturbation patterns as for the isolated C2 domains (27). These data indicate that Pb^2+^ selectively populates Site 1 of the C2A and C2B domains in the context of the C2AB fragment and that the linker region is not involved in metal-ion binding. The state of C2AB with Pb^2+^ bound to Site 1 in both domains is subsequently referred to as C2AB·Pb1.

Addition of anionic bicelles to the C2AB·Pb1 complex produced the following spectroscopic signatures in the [^15^N-^1^H] TROSY-HSQC spectra: decrease of the N-H cross-peak intensities and chemical shift perturbations, both of which are indicative of the protein-membrane binding process. The % intensity decrease was rather uniform within each domain, but the extent was different: 28% and 55% on average for the C2B and C2A domains, respectively. The pattern of chemical shift perturbations and their magnitude, small for the C2A domain loops and large for the LRC and loop 3 of the C2B domain (**Fig. 7A**), mirrored the data obtained for the individual domains (**Figs. 4B** and **5B**).

**Figure 7.**
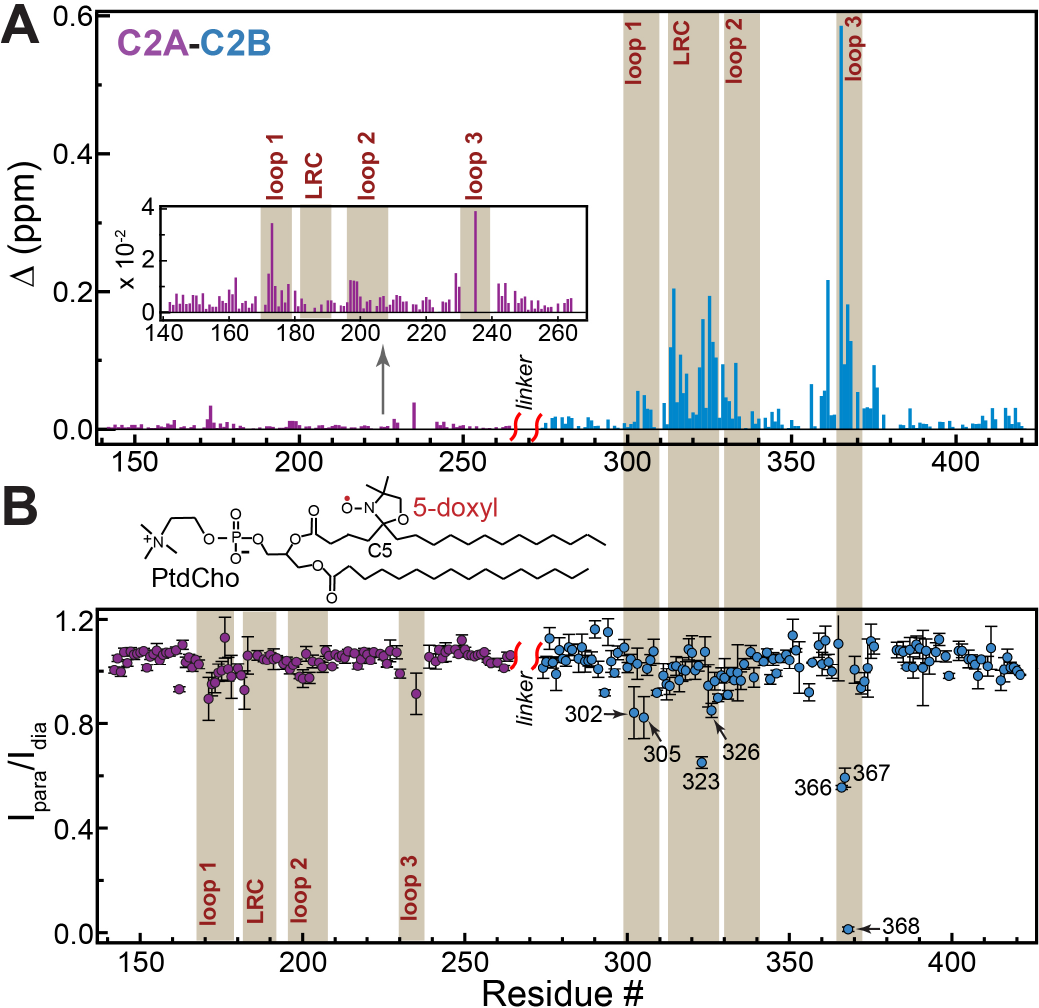
The C2AB·Pb1 complex binds to anionic bicelles with both domains contacting the membrane. (**A**) Loop 3 and LRC of the C2B domain make a dominant contribution to the chemical shift perturbation experienced by the C2AB·Pb1 complex upon binding anionic bicelles. The C2A data are shown in the inset with the re-scaled Y-axis. The CSP patterns of the C2 domains in the context of the C2AB fragment-bicelle system are similar to those of the individual domains shown in Figs. 4B and 5B. The similarities refer to both the overall pattern and the magnitude. (**B**) Membrane contact of C2A and C2B is revealed through the attenuation of the C2AB·Pb1 peak intensities upon addition of anionic bicelles containing a paramagnetic lipid, 5-doxyl PtdCho, at ~3 molecules per leaflet. The attenuation is expressed as residue-specific I_para_/I_dia_ ratios, where I_para_ and I_dia_ are the peak intensities in the presence and absence of paramagnetic lipids, respectively.

We used the paramagnetic relaxation enhancement (PRE) of NH groups to evaluate the extent of membrane contacts of C2 domains in the C2AB fragment. A lipid bearing a paramagnetic doxyl tag, 5- doxyl PC, was introduced into the bicelles to give ~3 molecules per leaflet (**Fig. 7B**). The unpaired electron of the doxyl group enhances the transverse relaxation rates of neighboring spins, resulting in broadening of the corresponding N-H cross-peaks and reduction in their intensities. The label position is such that N-H groups penetrating the headgroup region would experience the most broadening. If the protein-membrane binding process is in fast exchange on the NMR chemical shift timescale, which is the most likely scenario for the C2AB·Pb1 complex, the PRE effect is scaled by the population of paramagnetic species.

The residue-specific PRE values were calculated as the ratios of protein peak intensities in the C2AB·Pb1 samples containing diamagnetic and paramagnetic preparations of bicelles, I_dia_/I_para_ (**Fig. 7B**). Consistent with the chemical shift perturbation data of **Fig. 7A**, the C2A domain showed weak but readily discernable PRE effect in the loop regions. The PRE values were significantly larger for the C2B domain, with the residues of loop 3 (K366, I367, and G368) and LRC (L323 and K326) showing >15% intensity attenuation. Taken together, our data on the C2AB·Pb1 complex indicate that both domains make membrane contact when Site 1 on each domain is populated by a divalent metal ion. These results support the existence of a “primed” membrane-associated state under limiting metal ion conditions. In such a state, PtdIns(4,5)P_2_-mediated membrane docking of C2B will be augmented by the metal ion at Site 1 and keep both domains anchored at the membrane interface. The membrane binding of C2A will remain weak. This will ensure a strict metal-ion concentration dependence of the membrane insertion process for the C2A that requires a full complement of metal ions to associate with PtdSer-containing membranes.

### Progressive saturation of metal ion binding sites in full-length Syt1 results correlates with membrane penetration depth

To determine how the occupancy of metal ion binding sites influences the membrane insertion of the C2 domains in full-length Syt1, we conducted EPR experiments on the protein reconstituted into anionic LUVs. Under the conditions of our EPR experiments, both C2 domains of Syt1 bind in *cis* with respect to the transmembrane segment that anchors the protein to the LUVs (43). The spin-labeled sidechain R1 (**Fig. 8**) was attached to either loop 1 site in C2A (173R1) or a loop 1 site in C2B (304R1). Both sites are known to penetrate negatively charged membranes under saturating Ca^2+^ conditions. When no metal ions are present in solution, the R1 sidechain is in the aqueous phase and hence highly mobile, producing narrow EPR lineshapes for both C2 domains (**Fig. 8**, black traces). Addition of two-fold molar Pb^2+^ excess with respect to protein selectively populates Site 1 in C2A and C2B. The EPR spectra under these conditions (blue trace in **Fig. 8**) show significant broadening indicative of motional restriction due to partitioning of R1 into membranes. Addition of Pb^2+^ at a molar ratio (Pb^2+^:Syt1) of 20:1 gave rise to further broadening of the spectra, suggesting that binding of additional metal ions to the loop regions results in deeper membrane penetration of the C2 domains.

**Figure 8.**
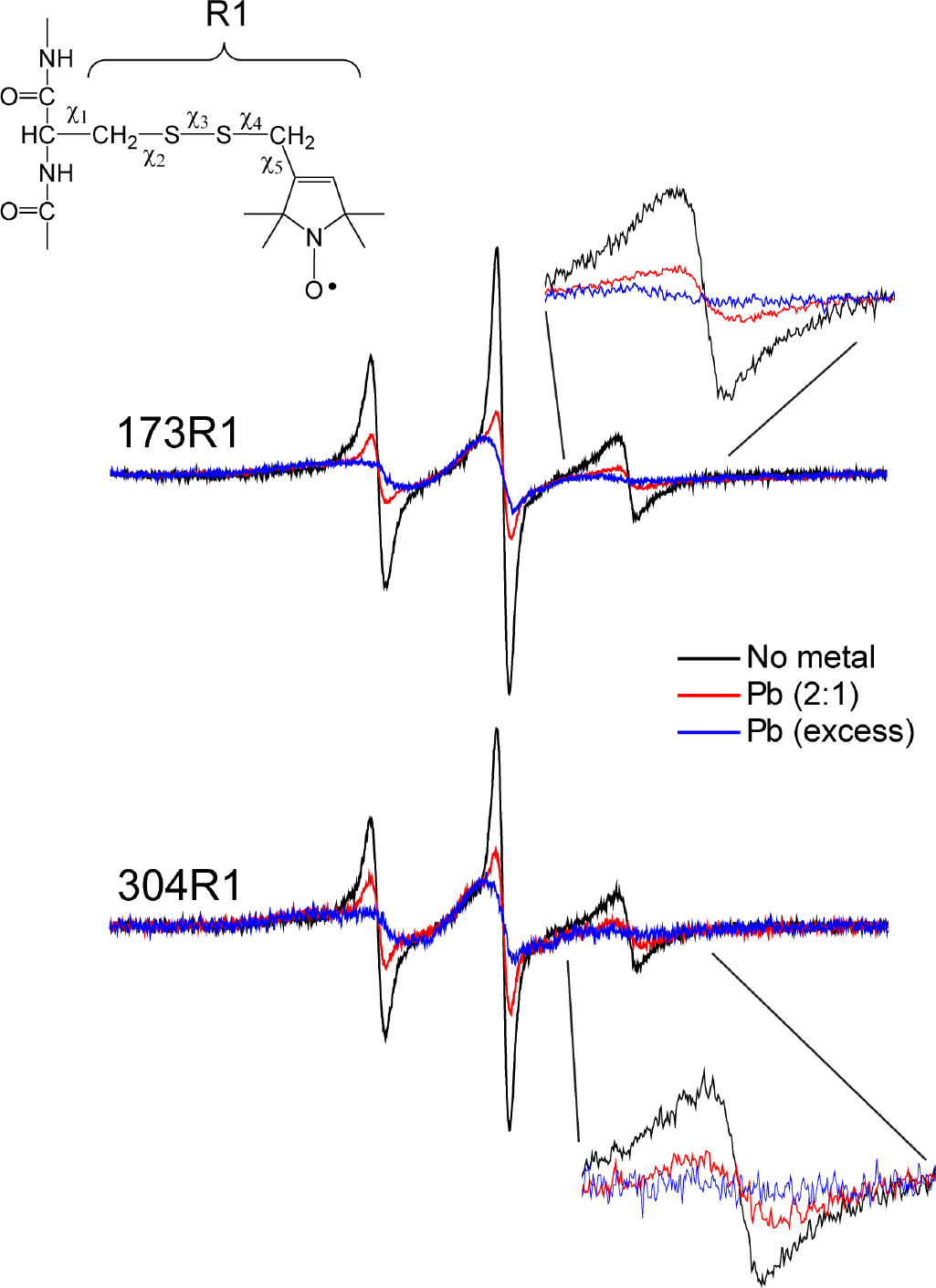
Membrane insertion of C2 domains in full-length Syt1 depends on the occupancy of metal-ion binding sites. X-band EPR spectra of full-length membrane-reconstituted Syt1, where the first Ca^2+^ binding loop of either the C2A domain (site 173) or the C2B domain (site 304) was labeled with MTSL to produce the R1 side-chain. Pb^2+^ binding to Syt1 promotes membrane insertion of the domains that is dependent upon the occupancy of the sites in the binding loops. The additional broadening in the EPR spectra with excess Pb^2+^ (Pb^2+^:Syt1, 20:1) suggests a deeper membrane penetration of the domains in this state. The ratio of Pb^2+^:Syt1 was varied from 12:1 to 50:1 with no change in the EPR spectra. The EPR spectra in the state labeled “No metal” were obtained after the addition of 4 mM EGTA, and these spectra were identical to spectra obtained prior to the addition of Pb^2+^.

To further validate our findings, we used progressive power saturation of the EPR spectra and a collision-gradient approach to determine the membrane depth of 173R1 and 304R1. **Table 1** gives the depth parameters and membrane depths under two Pb^2+^ concentration conditions. When Site 1 is occupied by a metal ion, both 173R1 (C2A) and 304R1 (C2B) are positioned on the aqueous side of the phosphate groups, with average distances of 0.8 Å and 3.5 Å, respectively, above the level of lipid phosphates. Binding a full complement of Pb^2+^ ions results in the R1 displacement into the membrane hydrocarbon region, with average distances of 3.6 Å (173R1, C2A) and 1.7 Å (304R1, C2B) below the phosphate level.

**Table 1.** Power saturation data for site 173R1 and 304R1when Syt1 is bound to membranes in the presence of Pb^2+^.

| Label | lipid | metal ion added<sup>§</sup> | depth parameter ($\Phi$) | Position ($\text{\AA}$)<sup>†</sup> |
| --- | --- | --- | --- | --- |
| FL SYT 173R1 | PC/PS (15%) | $\text{Pb}^{2+}$ 2:1 ($\text{Pb}^{2+}$:Syt1) | $-1.56 \pm 0.1$ | -0.78 |
| | | $\text{Pb}^{2+}$ excess | $-0.72 \pm 0.05$ | 3.59 |
| FL SYT 304R1 | PC/PS (15%) | $\text{Pb}^{2+}$ 2:1 ($\text{Pb}^{2+}$:Syt1) | $-1.75 \pm 0.1$ | -3.53 |
| | | $\text{Pb}^{2+}$ excess | $-1.19 \pm 0.1$ | 1.73 |

<sup>§</sup>Data for the 2:1 $\text{Pb}^{2+}$ to FL-Syt1 ratio (1:1 lead to C2 domain) were taken to examine membrane penetration for a partially bound lead state, where only Site1 is populated (27). Excess lead was then added to examine differences in membrane binding when both sites are populated in the C2 domains.
<sup>†</sup>The position indicates the average location of the nitroxide relative to the lipid phosphate plane. A negative value for the position indicates that the label is located on the aqueous side of the phosphate plane.

Both the EPR lineshapes and the membrane depth measurements support a more peripheral membrane-bound state of Syt1 when the metal ion binding sites have low occupancy. In the full-length protein the C2 domains are tethered to membranes via the N-terminal transmembrane region and this facilitates the formation of such a state by creating effectively high local lipid concentrations in the vicinity of the C2 domains. Furthermore, it “primes” Syt1 to further metal ion events by placing its C2 domains in close proximity to the negatively charged membrane interface.

## DISCUSSION

Ca^2+^-dependent membrane binding of Syt1 plays a crucial role in the excitation-secretion coupling by enabling the SNAREs to initiate vesicle fusion (7,47,48). Syt1-mediated evoked release has been detected at intracellular Ca^2+^ concentrations as low as 25 μM (49). However, Syt1 is an intrinsically weak Ca^2+^ sensor in the absence of anionic membranes, with affinities ranging between 50 μM to >10 mM for the 5 Ca^2+^-binding sites (17,18,50). In the vicinity of anionic phospholipids, the apparent Ca^2+^ affinity of Syt1 becomes ~3-4 μM (50). The anionic lipids play two roles: (i) mask the positively charged surface patches of C2 (e.g., the LRC) through transient interactions; and (ii) directly coordinate C2-bound divalent metal ions (31,32). It was originally hypothesized for the C2 domain of PKCα that the initial membrane association step requires its interactions with cytosolic Ca^2+^ through the highest-affinity metal ion-binding site (20). Pb^2+^, a xenobiotic metal ion that is a structural and functional surrogate of Ca^2+^ gave us access to these states of the Syt1 C2 domains in three protein contexts: isolated domains, the C2AB fragment, and full-length Syt1.

Apart from the clear changes in electrostatics, population of Site 1 by a divalent metal ion alters the dynamic behavior of the membrane-binding regions. Both C2 domains in the metal-free state have a highly dynamic loop 3 that undergoes conformational exchange on a timescale <100 μs (**Fig. 2**). The attenuation of loop 3 dynamics caused by a metal ion at Site 1 (**Fig. 3**) could facilitate the metal-ion dependent membrane-binding for the following three reasons: (i) loop 3 provides half of oxygen ligands of the metal ion at Site 1 and >60% of the coordinating ligands required for metal ion at Site 2; (ii) loop 3 harbors key positively charged residues (R233 in C2A; and K366 in C2B) (11,51,52) that can potentially interact with anionic lipid headgroups upon partial or full neutralization of the intra-loop region; and (iii) loop 3 has two hydrophobic residues (F231 and F234 for C2A; Y364 and I367 in C2B) that could potentially interact with the hydrophobic part of the bilayer (10,11,53). In addition to loop 3, the LRC region in apo C2A, but not apo C2B, shows conformational dynamics that gets attenuated by metal ion binding to Site 1. The allosteric nature of this effect is surprising, as the LRC is positioned >10 Å away from Site 1. The functional significance of these changes in C2A dynamics remains to be investigated.

With respect to the membrane-binding properties, metal ion at Site 1 enhances the membrane interactions of the C2 domains, individually and in the C2AB fragment (**Figs. 4**-**7**). In C2A, it serves to promote weak association with anionic membranes containing either PtdSer or both PtdSer and PtdIns(4,5)P_2_, in direct support of the hypothesis originally proposed for the C2-domain mediated recruitment of PKC to membranes (20). In C2B, the metal ion in Site 1 serves to strengthen its metal-ion independent association with PtdIns(4,5)P_2_ (**Fig. 6**). There was no obvious manifestation of avidity in our experiments on the C2AB fragment (**Fig. 7**), suggesting that either the linker region connecting the two domains is sufficiently flexible for the domains to behave independently, or the C2 domains bind in “trans” relative to each other.

The experiments on full-length Syt1, where the protein is tethered to the membrane via the N-terminal helical segment, provided the clearest picture of the effect that sequential population of metal-ion binding sites may have on the membrane interactions of the C2 domains. The EPR data (**Fig. 8** and **Table 1**) clearly show the correlation between the number of metal sites populated in the intra-loop region and the depth of membrane insertion. In the single metal-ion bound states, the loop 1 probes are above the lipid phosphate plane. Binding of additional metal ions positions the loop 1 probes below that plane, into the acyl chain region.

If we extrapolate our findings to the native metal ion Ca^2+^, our data support the existence of a shallow membrane-bound state of Syt1 that is formed at the initial stages of C2-membrane recruitment, when cytosolic Ca^2+^ interacts with the C2 domains prior to their binding to PtdSer (**Fig. 9**). In such a state, both domains make membrane contact: C2A through the loop regions, and C2B through its LRC region, with the latter interaction enhanced but not driven by the metal ion in Site 1. The conformational dynamics of the loops is significantly attenuated, and serves to pre-organize the loops for subsequent metal-ion and membrane binding events. This “primed” state, in combination with high local concentration of anionic lipids (54,55) and Ca^2+^(56,57) at the presynaptic membranes will lower the Ca^2+^ concentration requirements for full membrane insertion of Syt1. The proximity of the “primed state” to the anionic membrane interface would therefore turn the C2 domains, that lack cooperativity with respect to Ca^2+^ binding in solution, into highly cooperative modules that are capable of responding to physiological Ca^2+^ concentrations.

**Figure 9.**
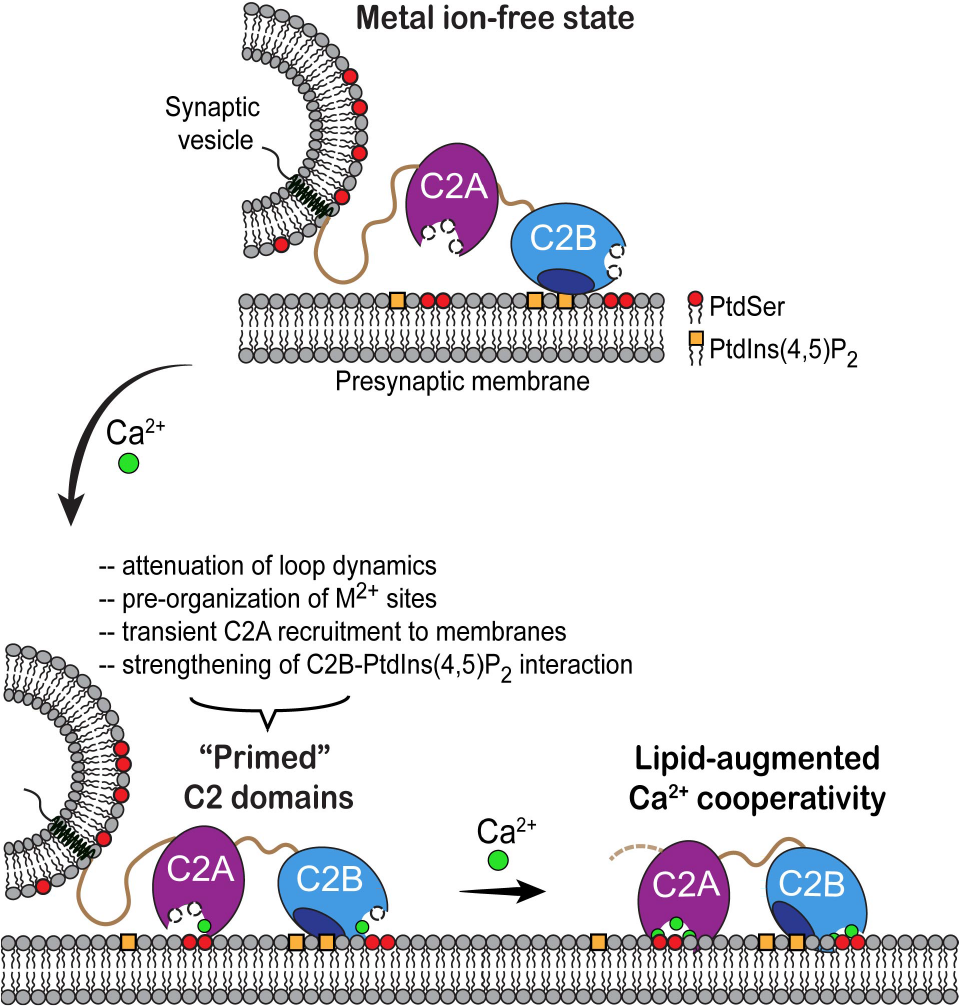
Priming of Syt1-membrane interactions by partial saturation of intra-loop metal-ion binding sites. In the metal ion-free state, only C2B domain is interacting with the PtdIns(4,5)P_2_ component of the presynaptic membrane; there are no interactions with PtdSer through the loop regions of either C2 domain (SNAREs or interactions of C2B domain with SNAREs are not depicted for simplicity). Binding of cytosolic Ca^2+^ to Site 1 will drive the association of C2A with the anionic lipids of the membrane and enhance the C2B-PtdIns(4,5)P_2_ interactions. Both C2 domains adopt a shallow membrane-bound state, in which the conformational flexibility of the loop regions (and the LRC region of C2A) is attenuated. The proximity of C2 domains to anionic phospholipids enables the cooperative binding of Ca^2+^ ions to the intra-loop region to achieve full saturation and membrane insertion.

## EXPERIMENTAL PROCEDURES

### Materials

The murine Syt1 cDNA was purchased from Open Biosystems (GE Life Sciences). Concentrated stock solutions of Pb^2+^ were prepared by dissolving Lead acetate tri-hydrate (Sigma-Aldrich) in HPLC-grade water or decalcified buffers. The necessary dilutions of this stock solution were freshly prepared prior to use to make working solutions. All the buffer solutions used in the experiments (MES/Bis-tris, Sigma-Aldrich) were treated with the ion-chelating resin Chelex-100 (Sigma-Aldrich) to remove trace divalent metals before use. Lipid components used in the phospholipid vesicle as well as bicelle preparations: 1-palmitoyl-2-oleoyl-sn-glycero-3-phosphocholine (POPC), 1-palmitoyl-2-oleoyl-sn-glycero-3-phospho-L-serine (POPS), 1,2-dimyristoyl-sn-glycero-3-phosphocholine (DMPC), 1,2-diheptanoyl-sn-glycero-3-phosphocholine (DHPC), 1,2-dimyristoyl-sn-glycero-3-phospho-L-serine (DMPS), L-α-phosphatidylinositol-4,5-bisphosphate (PtdIns(4,5)P_2_) were obtained from Avanti Polar Lipids Inc. (Alabaster, AL). The short-chain Phosphatidylinositol 4,5-bisphosphate (di-C4-PtdIns(4,5)P_2_) was obtained from Echelon Biosciences.

### Protein expression and purification

The gene segments encoding Syt1 C2A (residues 137-265), C2B (residues 271-421), and C2AB (residues 137-421) were cloned into pET-SUMO vector (Novagen, Madison, WI) and expressed as 6xHis-tagged SUMO fusion proteins in the *E. coli* BL21(DE3) (C2A, C2AB) and Rosetta(DE3) (C2B) strains as described previously (22,27). The full length-Syt1 (FL-Syt1) (residues 1-421) with the native cysteines mutated as: C73A, C74A, C76A, C78A, C82S, and C277S was cloned into a pET-28a vector. For EPR measurements, additional cysteine mutation were introduced into the cysteine-free FL-Syt1 at either positions M173C or V304C and expressed as 6xHis-tagged proteins in the *E. coli* BL21(DE3)-RIL strain as described before (22,43). All mutants were prepared using the QuickChange site-directed mutagenesis kit (Stratagene, La Jolla, CA.) and verified by DNA sequencing (Genewiz, South Plainfield, NJ).

Expression and purification steps from the previously described protocols were followed (22,27,43). Briefly, all SUMO fusion proteins were purified from cell lysates using HisTrap HP columns followed by removal of 6xHis-SUMO fusion tag using SUMO protease. The partially purified proteins were further refined using anion (HiTrap Q HP, C2A and C2AB) and cation (HiTrap SP HP, C2B)-exchange chromatography steps to remove any charged protein or nucleic acid impurities. The ion-exchange chromatography buffers were also supplemented with 100 μM EDTA to ensure that the purified Syt1 domains are free of metal contamination. Immediately prior to use, the protein stock solutions were concentrated and subjected to four successive passes through the desalting PD MidiTrap G-25 columns to attain complete buffer-exchange and removal of EDTA.

For the NMR measurements, individual C2 domains were uniformly enriched with ^15^N, as described previously (22,27). The C2AB fragment was additionally deuterated to a level of ~80%, by growing *E. coli* on the M9 medium supplemented with D_2_O and BioExpress^®^ 1000 (10 ml/L; [U-^2^H 98%, U-^15^N 98%]). The isotopically enriched chemicals were purchased from Cambridge Isotope Laboratories, Inc. Unless specified otherwise, all isotopically enriched protein preparations were exchanged post-purification into an NMR buffer of the composition: 20 mM MES (pH 6.0), 100 mM KCl, 8% D_2_O, and 0.02% NaN_3_.

FL-Syt1 used for the EPR experiments was purified in CHAPS (3-[(3-Cholamidopropyl)dimethylammonio]-1 propanesulfonate) by affinity chromatography using Ni-NTA agarose resin (Qiagen). The protein was spin labeled overnight at 4 °C using the thiol-specific spin label, MTSL (1-oxy-2,2,5,5-tetramethylpyrrolinyl-3-methyl methanethiosulfonate) while bound to the Ni-NTA column. The labeled protein was eluted followed by removal of the 6xHis tag by thrombin cleavage and further purified by cation-exchange chromatography (HiTrap SP HP).

### Preparation of phospholipid vesicles and bicelles

Large unilamellar vesicles (LUVs) of desired compositions were prepared by aliquoting the chloroform solutions of the POPC/POPS lipids, followed by extensive vacuum drying and extrusion in NMR buffer (Mini-Extruder, Avanti Polar Lipids, Inc.). All LUV preparations were verified for the mean diameter of 100 nm by dynamic light scattering and used within 2 days of preparation. Phospholipid concentrations in LUV solutions were calibrated using the phosphate determination assay (58).

To prepare isotropically tumbling bicelles, chloroform solutions of DMPC and DHPC were aliquoted, extensively dried under vacuum and resuspended in NMR buffer. DMPC preparation was vortexed and subjected to 3 rapid freeze-thaw cycles to create homogeneous slurry. Clear DHPC solution was then added to achieve two-fold molar excess to DMPC (q=0.5). The resulting mixture was briefly vortexed and subjected to 4 rapid freeze-thaw cycles, resulting in clear homogeneous bicelle stock solutions. Total lipid concentration was verified using phosphate determination assay (58). Additional long-chain lipid components, such as DMPS and PtdIns(4,5)P_2_, were dried and added during the bicelle preparation wherever applicable. The bicelle stock solutions were flash frozen and stored at −20 °C. Before use, the frozen stocks were thawed at 42 °C, followed by incubation at room temperature.

### Nuclear Magnetic Resonance (NMR) spectroscopy

All NMR experiments were conducted at 298.2 K, unless specified otherwise; the temperature was calibrated using deuterated methanol. NMR data were processed with NMRPipe (59) and analyzed with Sparky (60). Curvefit program (Arthur G. Palmer’s laboratory, Columbia University) was used to fit the relaxation data and extract relaxation parameters.

### Measurements of ^15^N transverse relaxation rate constants, R_2_

The R_2_ values were measured on an NMR spectrometer (Avance III HD, Bruker Biospin) operating at the ^1^H Larmor frequency of 600 MHz (14.1 T) and equipped with a cryogenically cooled probe. NMR samples contained 300 μM [U-^15^N]-enriched C2A or C2B, each in the absence or presence of stoichiometric amounts of Pb^2+^. Additional relaxation measurements were conducted on the Pb^2+^-complexed C2A domain in the presence of LUVs. The site-specific R_2,CPMG_ values (61) were extracted from fitting the ^15^N-^1^H_N_ cross-peak intensity decays at seven unique and three duplicate relaxation time points ranging from 8 to 140 ms. The free-precession R_2,HE_ values (62) were measured to estimate the exchange contribution to the observed relaxation behavior and the timescale of exchange, both through direct comparison with the R_2,CPMG_ values. The Hahn-Echo delay was set to 48.2 ms. All relaxation data were acquired in the interleaved manner.

### NMR-detected interactions between Syt1 domains and their ligands

#### C2AB and Pb^2+^

Binding of Pb^2+^ to C2AB was monitored using ^15^N-^1^H TROSY-HSQC spectra collected on an NMR spectrometer (Avance III HD, Bruker Biospin) operating at the ^1^H Larmor frequency of 800 MHz (18.8 T) and equipped with a cryogenically cooled probe. Pb^2+^ was added to the NMR sample containing 100 μM [U-^15^N, ~80%-^2^H] C2AB at 1- and 2-fold stoichiometric excess. The chemical shift perturbations (CSPs, Δ) due to Pb^2+^ interactions with proteins were calculated using the following equation:

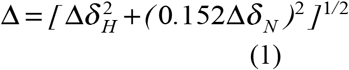

where Δδ_H_ and Δδ_N_ are residue-specific ^1^H and ^15^N chemical shift differences between metal-free and Pb^2+^-complexed states of C2AB.

#### Syt1 domains and bicelles

Binding of isotropic anionic bicelles to metal-free and Pb^2+^-complexed C2A, C2B and C2AB was conducted at 18.8 T and 300 K. A series of ^1^H-^15^N HSQC (TROSY-HSQC) spectra were acquired at each sample condition for the C2A/C2B (C2AB) domains. Protein concentration was kept at 150 μM. The bicelles were added to a total lipid concentration of 60 mM. The bilayer compositions of the added bicelles were either DMPC:DMPS:PtdIns(4,5)P_2_=84:14:2 or DMPC:DMPS=85:15. The chemical shift perturbations induced by bicelle interactions were calculated using eq. (1). The absolute resonance intensities for the bicelle-containing samples were normalized to their respective bicelle-free reference counterparts in order to estimate the signal attenuation.

#### C2A and LUVs

The interaction of the C2A domain with LUVs composed of either pure POPC or POPC:POPS=80:20 was monitored at 14.1 T using a 100 μM sample of [U-^15^N] C2A. A series of ^15^N-^1^H HSQC spectra were collected in the presence of LUVs, with the total lipid concentration ranging from 0 to 2.0 mM. Residue-specific intensity changes were distributed into linearly and exponentially decaying patterns based on their behavior, and the respective fitting models were used to extract the “decay” rates.

#### C2B and di-C4-PtdIns(4,5)P_2_

Interactions of C2B with di-C4-PtdIns(4,5)P_2_ were monitored at 14.1 T, using 100 μM C2B in either apo- or Pb^2+^- complexed forms. The di-C4-PtdIns(4,5)P_2_ concentrations were: 0, 10, 50, 100, 150, 300, and 550 μM for the C2B·Pb1 sample; and 55, 100, 150, 250, 400, 600, 800, and 1000 μM for the apo C2B sample. Residue-specific chemical shift perturbation values Δ were calculated using Eq. (1). The binding curves were constructed by plotting Δ as a function of di-C4-PtdIns(4,5)P_2_ concentration. To extract the dissociation constant *K_d_*, the curves were globally fitted (C2B·Pb1: 7 residues, apo C2B: 5 residues) with the single-site binding model:

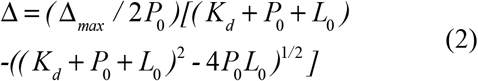

where Δ_*max*_ is the maximum perturbation, and *P*_*0*_ and *L*_*0*_ are the total protein and di-C4-PtdIns(4,5)P_2_ concentrations, respectively.

### Measurements of Electron Paramagnetic Resonance (EPR) spectra

For EPR measurements, FL-Syt1 was reconstituted into POPC:POPS=85:15 at a 1:200 protein to lipid ratio and dialyzed into metal-free buffer (20 mM HEPES, 150 mM KCl, pH 7.4) in the presence of Bio-Beads (Bio-Rad Laboratories, Hercules, CA) (43). Pb^2+^ was then added to samples at either a 2:1 Pb^2+^:FL-Syt1 ratio or in 12-to 50-fold excess. Measurements were taken as described previously, using a Bruker X-Band EMX spectrometer (Bruker Biospin, Billerica, MA) equipped with an ER 4123D dielectric resonator. Continuous wave spectra were taken with 100 G magnetic field sweep, 1 G modulation, and 2.0-milliwatt incident microwave power at room temperature. 10 μL samples of protein with concentrations varying between 15-75 μM depending on the desired Pb^2+^:FL-Syt1 ratio were prepared in glass capillary tubes (0.60 mm inner diameter × 0.84 mm outer diameter round capillary; VitroCom, Mountain Lakes, NJ). The phasing, normalization, and subtraction of EPR spectra were performed using in-Lab software written by David Nyenhuis and LabVIEW software provided by Dr. Christian Altenbach (43). Progressive power saturation of the EPR spectrum was used to determine nitroxide membrane depth and was performed as previously described (22,43). Samples were run in gas-permeable TPX-2 capillaries, and the values of Δ*P*_1/2_ obtained in air and in the presence of Ni(II)EDDA were used to calculate a depth parameter, Φ (63). The spin label depth was then estimated using the empirical expression:

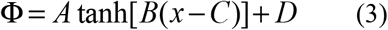

where *x* is the distance of the spin label from the lipid phosphate plane (positive *x* values are inside while the negative values are outside the bilayer) and *A*, *B*, *C*, and *D* are empirically determined constants (22,43).

## Author contributions

T.I.I., D.S.C., S.K., and S.B.N. designed the study; S.K., S.B.N., and B.H. conducted the experiments; S.K. and S.B.N. analyzed the data; T.I.I., S.K., D.S.C., and S.B.N. wrote the manuscript.

## The abbreviations used are

Syt1: Synaptotagmin 1
PtdSer: phosphatidylserine
PtdCho: phosphatidylcholine
PtdIns(4,5)P_2_: phosphatidylinositol 4,5-bisphosphate
SNAREs: soluble N-ethylmaleimide sensitive factor attachment protein receptors
LRC: lysine rich cluster
CPMG: Carr-Purcell-Meiboom-Gill
HE: Hahn-Echo
HSQC: heteronuclear single quantum coherence
TROSY: transverse relaxation optimized spectroscopy
CSPs: chemical shift perturbations
PRE: paramagnetic relaxation enhancement

