## SUPPORTING INFORMATION for "Partial metal ion saturation of C2 domains primes Syt1-membrane interactions"

Running title: *Priming of Syt1-membrane interactions*

*To whom correspondence should be addressed:
Tatyana I. Igumenova: Department of Biochemistry and Biophysics, Texas A&M University, 300 Olsen Boulevard, College Station, TX 77843, United States;; Tel. (979) 845-6312

**MATERIAL INCLUDED: SUPPORTING FIGURES S1-S7**

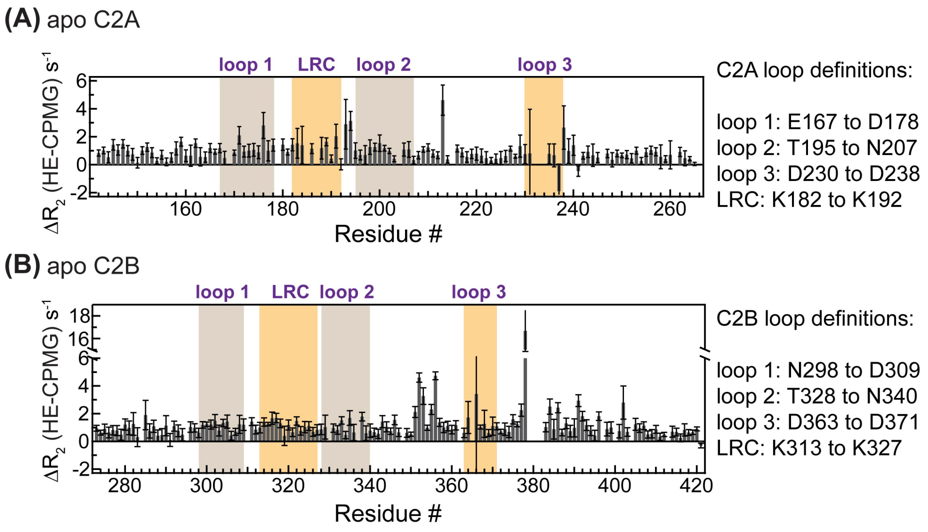


**Figure S1.** Small differences between residue-specific $R_{2,HE}$ and $R_{2,CPMG}$ values indicate that the timescale of the membrane-binding region dynamics in apo C2A (**A**) and C2B (**B**) domains is faster than 100 μs. This figure represents difference plots of the data shown in Figure 2.

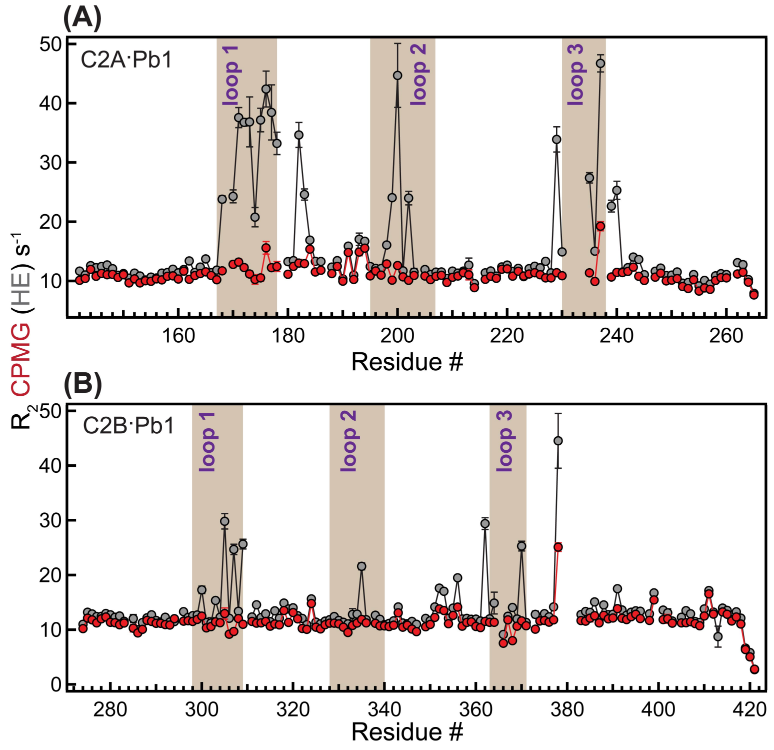


**Figure S2. Evidence for the millisecond (ms)-timescale dynamics associated with under-population of Site 1 by $Pb^{2+}$ in C2A (A) and C2B (B).** The conclusion regarding the timescale comes from the comparison of the residue-specific transverse relaxation rate constants $R_2$ obtained in the Hahn-Echo (HE, gray) and Carr-Purcell-Meiboom-Gill (CPMG, red) experiments. Application of the CPMG pulse train results in almost complete attenuation of chemical exchange contributions to the $R_{2,CPMG}$ values, which is an unambiguous evidence for the ms-timescale dynamics.

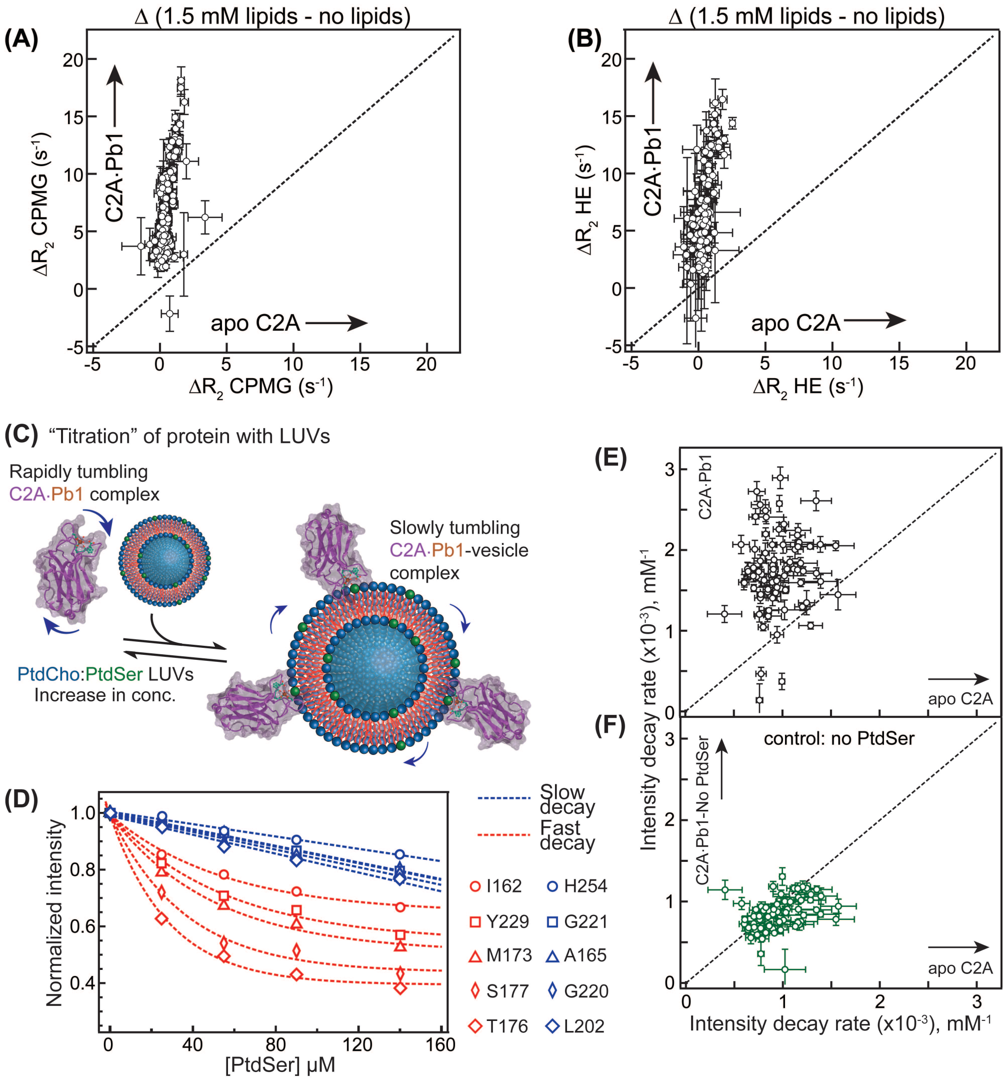


**Figure S3. Syt1 C2A·Pb1 complex binds to PtdSer-containing large unilamellar vesicles (LUVs).** (**A,B**) Correlation plots of $\Delta R_{2,CPMG}$ (**A**) and $\Delta R_{2,HE}$ (**B**) values in the C2A·Pb1 complex and apo C2A. $\Delta R_{2,CPMG}$ and $\Delta R_{2,HE}$ are calculated as the differences in residue-specific $R_{2,CPMG}$ and $R_{2,HE}$ values, respectively, in the presence and absence of LUVs. While no significant changes are observed in apo C2A upon addition of LUVs, all $\Delta R_{2,CPMG}$ and $\Delta R_{2,HE}$ values are > 0 in the C2A·Pb1 complex. These data indicate that PtdSer as anionic lipid component is necessary and sufficient for C2A to interact with membrane mimics upon binding a single metal ion to Site 1. (**C**) Schematic representation of the experimental design to probe the interaction of C2A·Pb1 with LUVs. The experiments were carried out in the "titration" mode, where increasing concentrations of LUVs were added to the [U-$^{15}$N] enriched C2A·Pb1 complex. The $^{15}$N-$^{1}$H HSQC spectra of C2A·Pb1 were recorded at each LUV concentration point. (**D**) Normalized N-H cross-peak intensities of C2A·Pb1 show consistent decay upon increase of LUV concentration, plotted along the X-axis as accessible PtdSer. Two types of behavior were observed, depending on the chemical exchange regime of a given residue, slow (blue) and fast (red). (**E,F**) Correlation plots of the intensity decay rates between C2A·Pb1 and apo C2A. In the presence of PtdSer-containing LUVs, the intensity decay rates of the C2A·Pb1 exceed those of apo C2A due to the interactions of the former with LUVs. This behavior is dependent on the presence of PtdSer in the membrane: omission of PtdSer results in comparable intensity decay rates that are mostly due to the increase in sample viscosity in both protein samples.

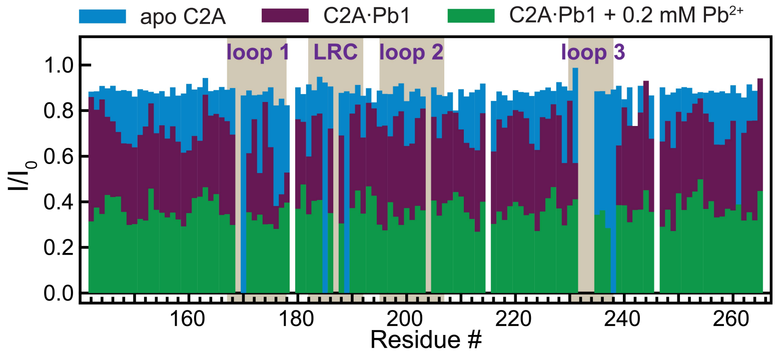


**Figure S4. Progressive population of metal ion binding sites by $Pb^{2+}$ in C2A drives the protein association with anionic membranes.** The ratio of the C2A N-H cross-peak intensities in the absence ($I_0$) and presence (I) of PtdSer-containing LUVs is plotted against the primary structure for three protein samples in different states of metal ligation. The composition of LUVs is PtdCho:PtdSer=80:20; the protein-to-lipid ratios are 1:20, 1:14, and 1:10 for the apo, C2A·Pb1, and (C2A·Pb1+0.2 mM $Pb^{2+}$) samples, respectively.

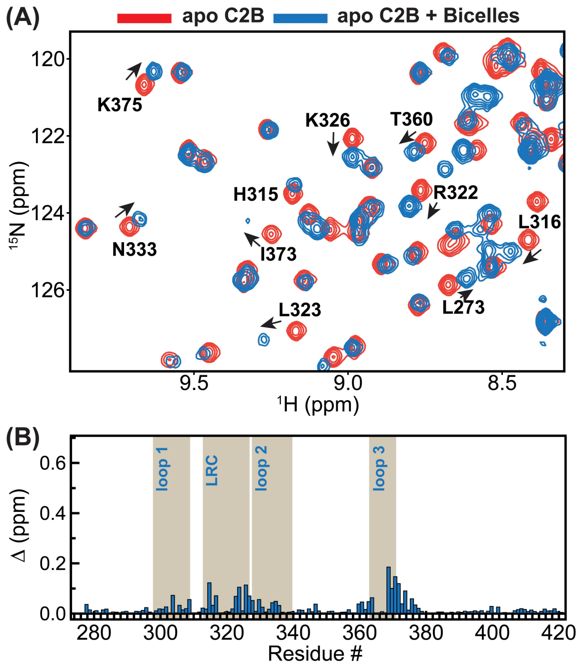


**Figure S5. Apo C2B binds to anionic bicelles that contain PtdIns(4,5)$P_2$ component.** (**A**) Overlay of the $^{15}$N-$^{1}$H HSQC spectral regions of apo C2B in the absence (red) and presence (blue) of anionic bicelles that illustrates CSPs experienced by the protein upon interactions with bicelles. The bicelle-containing spectrum is acquired with twice the number of scans compared to the bicelle-free one and depicted at the same contour threshold. (**B**) CSP plot calculated for the bicelle-containing C2B sample relative to the bicelle-free protein. The Y-axis scale is identical to that of the C2B·Pb1 plot shown in **Fig 5B**. The concentrations of protein and bicelle-lipids were 150 μM and 60 mM (~230 μM bicelles) respectively.

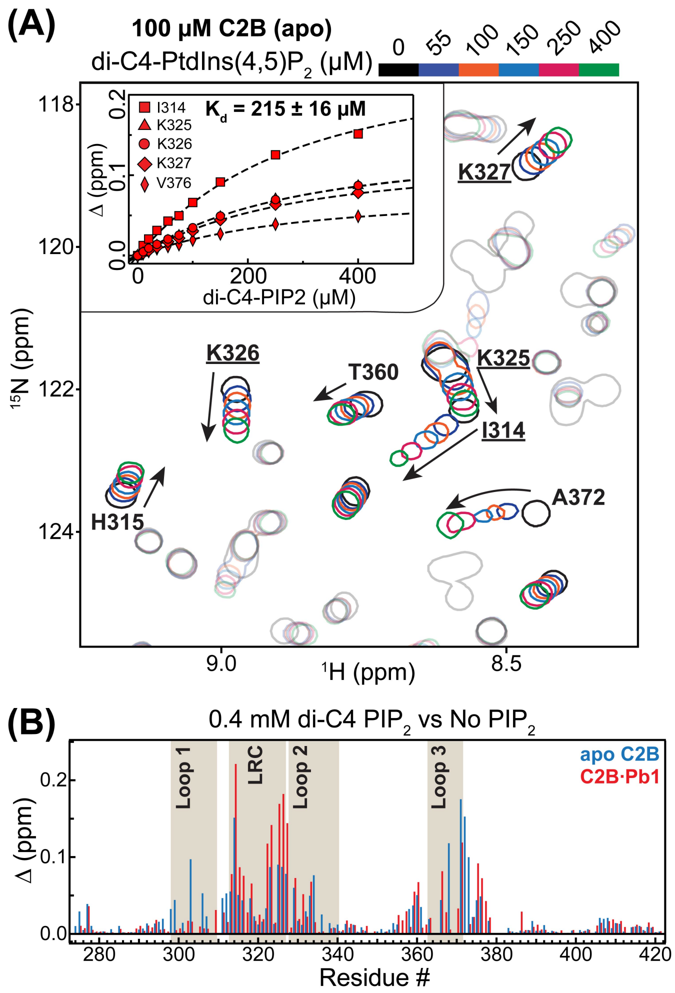


**Figure S6. PtdIns(4,5)$P_2$ binds to apo C2B with 2-fold lower affinity than to C2A·Pb1.** (**A**) Overlay of the [$^{15}$N-$^{1}$H] HSQC spectral region showing progressive chemical shift changes upon addition of di-C4-PtdIns(4,5)$P_2$ to apo C2B. N-H cross-peaks that illustrate the binding of di-C4-PtdIns(4,5)$P_2$ to the canonical site in the LRC are labeled. Inset: NMR-detected binding curves for residues that either form or are in close proximity to the canonical binding site. The effective $K_d$ value extracted from the global fit of the curves is 215 μM. (**B**) Comparison of the chemical shift perturbations Δ due to di-C4-PtdIns(4,5)$P_2$ binding (0-0.4 mM range) for apo C2B (blue) and the C2B.Pb1 complex (red).

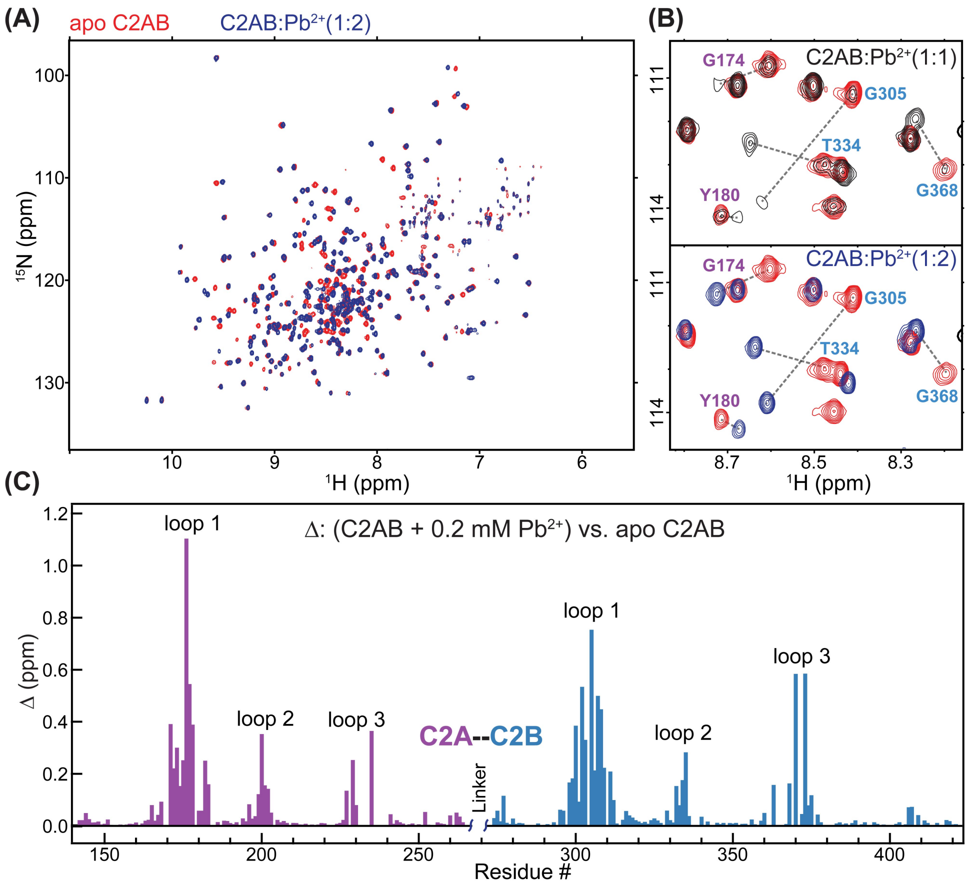


**Figure S7. Syt1 C2AB binds one $Pb^{2+}$ per C2 domain under C2AB:$Pb^{2+}$=1:2 conditions.** (A) Overlay of the [$^{15}$N-$^{1}$H] TROSY-HSQC spectra of apo (red) and $Pb^{2+}$-complexed (blue) C2AB showing the chemical shift changes upon $Pb^{2+}$ binding to the C2AB domain. The concentrations of C2AB and $Pb^{2+}$ were 100 and 200 µM, respectively. (B) Expansion of the [$^{15}$N-$^{1}$H] TROSY-HSQC spectra at protein:$Pb^{2+}$ =1:1 ratio (top) show that loop region residues are in slow exchange regime on the NMR chemical shift timescale. This is manifested in two sets of N-H cross-peaks: one set corresponding to the apo C2AB, and the other set corresponding to the $Pb^{2+}$-complexed C2AB; the peaks that belong to the same residue are connected with a dashed line. The N-H cross-peaks corresponding to the C2A and C2B residues are labeled in purple and blue, respectively. The spectrum of apo C2AB (red) is overlaid onto the spectrum of the $Pb^{2+}$-complexed C2AB for comparison. Addition of $Pb^{2+}$ to a final protein-to-$Pb^{2+}$ ratio of 1:2 results in complete redistribution of peak intensities towards the $Pb^{2+}$-complexed form (bottom). (C) Chemical shift perturbation plot showing that $Pb^{2+}$ binding affects the loop regions of both C2A and C2B. The pattern is consistent with that observed for the individual C2A and C2B domains with $Pb^{2+}$ bound to Site 1 (Katti, S., Her, B., Srivastava, A. K., Taylor, A. B., Lockless, S. W., and Igumenova, T. I. (2018) High affinity interactions of $Pb^{2+}$ with synaptotagmin I. Metallomics 10, 1211-1222).
